# Weakened airway epithelial junctions and enhanced neutrophil elastase release contribute to age-dependent bacteremia risk following pneumococcal pneumonia

**DOI:** 10.1101/2024.10.06.616883

**Authors:** Shuying Xu, Tianmou Zhu, Hongmei Mou, Shumin Tan, John M. Leong

**Affiliations:** Department of Molecular Biology and Microbiology, Tufts University School of Medicine, Boston, MA; Graduate Program in Immunology, Tufts Graduate School of Biomedical Sciences, Boston, MA; Mucosal Immunology and Biology Research Center, Massachusetts General Hospital, Boston, MA; Stuart B Levy Center for the Integrated Management of Antimicrobial Resistance, Tufts University, Boston, MA

**Author notes:** Corresponding author: John M. Leong.

**Keywords:** *Streptococcus pneumoniae*, pneumolysin, E-cadherin, neutrophil transmigration, aging

## Abstract

*Streptococcus pneumoniae* (*Sp*; pneumococcus), the most common agent of community-acquired pneumonia, can spread systemically, particularly in the elderly, highlighting the need for adjunctive therapies. The airway epithelial barrier defends against bacteremia and is dependent upon apical junctional complex (AJC) proteins such as E-cadherin. After mouse lung challenge, pneumolysin (PLY), a key *Sp* virulence factor, stimulates epithelial secretion of an inflammatory eicosanoid, triggering the infiltration of polymorphonuclear leukocytes (PMNs) that secrete high levels of neutrophil elastase (NE), thus promoting epithelial damage and systemic infection. Here, pulmonary E-cadherin staining of intratracheally inoculated mice revealed PLY-mediated disruption of AJC independently of PMNs. Apical infection of air-liquid interface (ALI) respiratory epithelial monolayers similarly showed that PLY disrupts AJCs. This epithelial damage promoted PMN transmigration and bacterial apical-to-basolateral translocation, and pharmacologically fortifying epithelial barrier function diminished both barrier breach *in vitro* and bacteremia *in vivo*. E-cadherin staining after *Sp* intratracheal inoculation of >20-month-old mice, or apical infection of ALI monolayers derived from these mice, revealed an age-associated vulnerability to PLY-mediated AJC disruption, which in turn enhanced PMN migration and bacteremia. In addition, we found that PMNs from aged mice secrete increased levels of tissue-damaging NE. Simultaneous pharmacological inhibition of tissue-destructive NE and fortification of pulmonary epithelial barrier function was required to reduce the level of *Sp* bacteremia in aged mice to that of young mice. This work underscores the importance of fully characterizing the multifactorial sources of age-associated susceptibility in devising adjunctive therapies to mitigate invasive pneumococcal disease in the elderly.

## INTRODUCTION

*Streptococcus pneumoniae* (*Sp*; pneumococcus) is a common asymptomatic colonizer of the nasopharynx but can cause serious infections including pneumonia, septicemia, and meningitis (Brown, Millett, Quint, & Orihuela, 2015), particularly in the elderly population (GBD Collaborators, 2018). Despite the availability of vaccines and antibiotics, *Sp* causes over 1 million deaths annually, most of which occur in individuals >65 years of age (GBD Collaborators, 2018). Hence, complementary treatment approaches targeting detrimental age-related changes in host response to *Sp* infection are needed, particularly those that limit complications highly associated with elderly individuals (Sundaresh et al., 2021).

A first line barrier against *Sp* spread is the integrity of the respiratory epithelium. Damage to the epithelium, as well as other tissues, is associated with excessive and/or sustained PMN presence in the airways (Ballinger & Standiford, 2010; T. Burns, Abadi, & Pirofski, 2005; Marks et al., 2007). The pore-forming toxin pneumolysin (PLY) is a key *Sp* virulence factor that promotes tissue damage and invasive disease. We previously found that apical infection of a polarized epithelial monolayer by PLY-producing *Sp* triggers 12-lipoxygenase activity and production of the eicosanoid hepoxilin A3 (HXA_3_), a potent PMN chemoattractant that both promotes PMN transepithelial migration and the release of neutrophil elastase (NE) (Xu et al., 2024), a protease with tissue-destructive potential (Boxio et al., 2016; Domon & Terao, 2021; Ginzberg et al., 2001). These PLY-induce transmigration of tissue-damaging PMNs disrupts the epithelial barrier, and inhibiting HXA_3_ production or NE activity diminishes lethal bacteremia following lung infection (Bhowmick, Clark, Bonventre, Leong, & McCormick, 2017; Bhowmick et al., 2013).

The detrimental effects of the acute inflammatory response to *Sp* lung infection are likely to be enhanced with age. In addition to age-associated functional defects in PMN functions, such as chemotactic accuracy (Sapey et al., 2017), phagocytosis (Simell et al., 2011), and ROS production (Biasi et al., 1996), the aged host experiences inflammaging, the elevation of inflammatory signaling (Frasca & Blomberg, 2016; Wenisch, Patruta, Daxböck, Krause, & Hörl, 2000). Indeed, *Sp-*infected elderly patients experience higher levels of PMN pulmonary influx (Menter, Giefing-Kroell, Grubeck-Loebenstein, & Tzankov, 2014; Pignatti, Ragnoli, Radaeli, Moscato, & Malerba, 2011), which is associated with increased bacterial burden and mortality.

These findings notwithstanding, the age-associated vulnerability to systemic *Sp* infection is complex and unlikely to be due solely to the enhanced epithelial disruption by PMNs. Epithelial barrier function is fostered by apical junctional complexes (AJC), comprised of tight junctions (TJ) and adherens junctions (AJ) (Ganesan, Comstock, & Sajjan, 2013). TJ proteins, such as zonula occludens (ZO), claudins, occludins, and junctional adhesion molecules (JAM), localize to the apical region of cell-cell junctions (Peter et al., 2017) and form an intercellular membrane fence (Otani & Furuse, 2020). AJ proteins, such as E-cadherin and catenins, are located basolateral to TJs and are essential for the formation and maturation of cell-cell contact (Bhatt, Rizvi, Batta, Kataria, & Jamora, 2013). Orchestration of the abundance and localization of junction proteins, as well as their interactions with the cytoskeleton and signaling modules, is essential for maintaining optimal epithelial barrier function (Ganesan et al., 2013). Loss of junctional integrity is associated with barrier permeability changes, and AJC malfunction of the pulmonary barrier often has deleterious outcomes during diseases of the lung, including pneumonia, acute lung injury, asthma, and chronic obstructive pulmonary disease (COPD) (Devaux, Mezouar, & Mege, 2019; Ghosh et al., 2022; Nawijn, Hackett, Postma, van Oosterhout, & Heijink, 2011; Wittekindt, 2017).

During pulmonary infection, *Sp* targets AJC proteins for disruption and compromises function of the airway epithelial barrier (Clarke, Francella, Huegel, & Weiser, 2011; LeMessurier, Hacker, Chi, Tuomanen, & Redecke, 2013; Peter et al., 2017). Pneumococcal infection reduces alveolar ZO-1, occludins, claudins, and cadherins in human lung explants (Peter et al., 2017) and murine models (Clarke et al., 2011; Cui et al., 2024; Jacques et al., 2020; Mo, Xu, Rosa, Hasan, & Adams, 2022). In addition, mice deficient in type I interferon, which are highly susceptible to severe bacteremia following lung challenge, show junctional defects in response to *Sp* infection (LeMessurier et al., 2013). Significantly, many genes associated with the assembly of AJCs are amongst those with the largest age-associated decrease in expression in the lungs (de Vries et al., 2017). Among them is *CDH1*, which encodes E-cadherin, a key regulator of epithelial barrier function (de Vries et al., 2022; Ghosh et al., 2022; Yuksel, Ocalan, & Yilmaz, 2021). E-cadherin dysfunction promotes cell turnover and mucin accumulation, contributing to loss of barrier integrity in asthma and COPD (Ghosh et al., 2022; Kim, Schein, & Nadel, 2005). Notably, we have reported that PLY promotes E-cadherin cleavage and dissolution during *Sp* apical infection of a monolayer of immortalized respiratory cells (Xu et al., 2023). However, whether age-related defects in AJC integrity contribute to susceptibility to *Sp* cross-epithelial dissemination or are further exacerbated by PLY remain to be elucidated.

In this study, we intratracheally challenged young or aged mice with *Sp*, assessing AJC disruption of the lung epithelium and subsequent bacteremia. Use of respiratory stem cell-derived air-liquid interface (ALI) epithelial monolayers derived from young or aged mice permitted *in vitro* assessment of age-related changes in epithelial AJC organization and barrier function. Additionally, ALI from mice that lack 12-lipoxygenase activity, which are thus incapable of HXA_3_ production, enabled documentation of a role of PLY-induced AJC disruption in *Sp*-promoted changes in barrier function independent of PLY-induced PMN chemoattractant secretion. Compared to lung epithelium from young mice, epithelium from aged mice was more susceptible to damage by PLY-producing *Sp*, resulting in greater PMN transmigration, barrier disruption, and bacterial translocation. Finally, pharmacologically counteracting AJC disruption mitigated *Sp*-induced epithelial damage and bacteremia in both young and aged mice; however, in aged mice, only the combined pharmacologic mitigation of PLY-induced changes to both PMN behavior and epithelial AJC integrity reduced systemic spread to that of untreated young mice.

## MATERIALS AND METHODS

### Bacterial strains and growth conditions

*Sp* TIGR4 (serotype 4) and the TIGR4 pneumolysin-deficient mutant (Δ*ply*) were a gift from Dr. Andrew Camilli (Tufts University School of Medicine, MA). Bacteria were grown to mid-exponential phase at 37 C at 5% CO_2_ in Todd Hewitt broth (BD Biosciences) supplemented with 0.5% yeast extract and Oxyrase (Oxyrase, Mansfield, OH), and frozen in growth media with 20% (v/v) glycerol. Bacterial titers in aliquots were confirmed by plating serial dilutions on Tryptic Soy Agar plates supplemented with 5% sheep blood (blood agar) (Northeast Laboratory Services, Winslow, ME). For experiments, frozen aliquots were grown in liquid culture and used at mid-log to late log phase.

### Murine infections

Young (2-month-old) BALB/c, C57BL/6J, and *Alox12/15* knockout (*Alox15*^*−/−*^) (B6.129S2-*ALOX15*^tm1Fun^/J) mice were obtained from Jackson Laboratories. Aged (>20-month-old) BALB/c and C57BL/6J mice were obtained from the National Institute on Aging aged rodent colonies. All animal experiments were performed in accordance with Tufts University Animal Care and Use Committee approved protocols. Bedding transfers to minimize microbiota differences between mice of different breeding facilities were performed on cages within the same mouse strain. Roughly one-quarter of soiled bedding was collected from each cage and the bedding from all cages were mixed in an empty sterile cage before redistribution across all cages.

To induce experimental pneumococcal pneumonia, young or aged BALB/c mice were intratracheally challenged with 1×10^7^ colony forming units (CFU) of *Sp* in 50 μl phosphate-buffered saline (PBS). Control mice received PBS. The role of preserving junctional integrity during on *Sp* infection was investigated by injection of either the Nrf2 agonist bardoxolone methyl (CDDO) at 100 μg/mouse, in 3% DMSO, 3% cremaphor EL (CrEL) in PBS alone, or in combination with the neutrophil elastase inhibitor Sivelestat at 500 μg/mouse, in PBS, intraperitoneally (*i*.*p*.) 1 hour prior to infection. Mice were euthanized at 18 h.p.i.. Blood was obtained by cardiac puncture. Bronchoalveolar lavage fluid (BALF) was collected by washing the lungs with 1 ml PBS via a cannula. Whole lungs were then removed, and bacterial burden enumerated by plating lung homogenate on blood agar plates.

### Lung barrier integrity by dextran and microscopy

Mice were intravenously injected with 70 kDa MW FITC-dextran at 5□mg/kg 30□minutes prior to euthanasia to assess lung permeability (Xu et al., 2024). Briefly, homogenized lungs were quantitated for FITC fluorescence using a Synergy H1 plate reader (BioTek) and readout was normalized to fluorescence in the serum of the same animal. To visualize airway epithelial junctions by fluorescence microscopy, lung tissues were harvested from euthanized mice, fixed in 4% paraformaldehyde, and sectioned to a thickness of 250 μm with a Leica Vibratome (0.145 mm/s, 70 Hz, blade angle 5°). Tissue sections were permeabilized with 0.1% Triton X-100 in PBS plus 3% bovine serum albumin (BSA) for 2 hours. Permeabilized sections were stained with anti-E-cadherin (24E10, Cell Signaling) and anti-Ly6G (1A8, BD Pharmingen) antibodies overnight, followed by Alexa Fluor 514-conjugated anti-rabbit and Alexa Fluor 647-conjugated anti-rat secondary antibodies, along with DAPI and Alexa Fluor 594-conjugated phalloidin (Invitrogen). Samples were mounted with Vectashield antifade mounting medium (Vector Laboratories) and visualized by confocal microscopy (Leica SP8).

### Measurement of PMN infiltration *in vivo*

For flow cytometric quantitation of lung PMNs, lung tissues were digested into a single cell suspension as previously described (Xu et al., 2024). Cells were resuspended in cell staining buffer (Biolegend) and stained on ice for 30 minutes with APC-conjugated anti-Ly6G (clone 1A8, Biolegend) and then washed two times in cell staining buffer (Biolegend). Cells were analyzed using a FACSCalibur flow cytometer (BD Biosciences) and the fluorescence intensities of the stained cells determined. Collected data were analyzed using FlowJo software (v10.7, BD) to determine the numbers of infiltrating (Ly6G^+^) PMNs.

### Establishment of epithelial air-liquid interface monolayers

Using a previously published airway basal cell isolation and expansion protocol (Gonzalez-Juarbe et al., 2017; Xu et al., 2024), healthy human donor-derived bronchial basal cells, young and aged C57BL/6J mouse-derived tracheal basal cells, and *Alox15*^−/−^ mouse tracheal basal cells were cultured in modified complete small airway epithelial growth media (SAGM) (Lonza, Cat. CC-3118)(Mou et al., 2016). Cells isolated from a single donor were used between passages 2-5 for consistency. To generate conventional upright monolayers on Transwells (Mou et al., 2016) for infection and imaging studies, the up-facing side of 6.5 mm Transwell inserts with 0.4 μm pores (Corning product #3470) were pre-coated with 804 G rat bladder cell conditioned medium as a source of collagen before cell seeding. For studying neutrophil transepithelial migration, the inverted ALI model was adopted (Yonker et al., 2017), where Transwells with permeable (3 μm pore size) polycarbonate membrane inserts (Corning #3415) were used and the underside of the Transwells were 804 G medium-coated before cell seeding. As previously described (Xu et al., 2024), each Transwell was seeded with 80 μl of airway basal cell suspension at a density of > 6000 cells/mm^2^, and cultured at air-liquid interphase in Pneumacult-ALI medium (StemCell Technology, Cat. 05001) for at least 21 days to allow for full epithelial maturation (Levardon, Yonker, Hurley, & Mou, 2018). Transepithelial electrical resistance was assessed using a voltmeter (EVOM2, Epithelial Voltohmmeter, World Precision Instruments, Inc.) to ensure the establishment of a polarized epithelial barrier.

### Infection of ALI monolayers

Apical surface of the ALI monolayers were infected with *Sp* at 1×10^7^ CFU in 25 μl of Hanks’ balanced salt solution (HBSS) supplemented with 1.2 mM Ca^2+^ and 0.5 mM Mg^2+^, and incubated at 37°C with 5% CO_2_ for 2 hours to allow for attachment and infection of the ALI monolayers. After treatment, Transwells were placed in 24-well receiving plates containing HBSS with Ca^2+^ and Mg^2+^, to allow for bacteria translocation for an additional 2 hours with or without the addition of 1×10^6^ PMNs to the basolateral chamber. 3,3′, 5,5′ tetramethylbenzidine dihydrochloride (TMB) peroxidase substrate conversion was used to detect flux of basally added horseradish peroxidase (HRP) to the apical chamber, as an assessment of ALI monolayer barrier integrity post-treatment. Buffer in the basolateral chambers was sampled and bacterial translocation across ALI monolayers was evaluated by plating serial dilutions on blood agar plates.

### PMN transepithelial migration assays

Whole blood obtained from healthy human volunteers under an IRB-approved protocol (Tufts University protocol #10489) was used to isolate neutrophils using the Easysep direct human neutrophil isolation kit (Stemcell). 1×10^6^ PMNs were added to the basolateral chamber after two hours of apical infection of the ALI monolayers with *Sp*.

After two hours of transmigration, PMNs in the apical chamber were quantified by MPO activity assay (Adams et al., 2019). Briefly, transmigrated PMNs were lysed by adding 50 μl of 10% Triton X-100 and 50 μl of 1 M citrate buffer and lysate was transferred to a 96-well plate. 100 μl of freshly prepared 2,2’-azinobis-3-ethylbenzotiazoline-6-sulfonic acid (ABTS) with hydrogen peroxide solution was added to each well and incubated in the dark at room temperature for 5-10 minutes. Absorbance at a wavelength of 405 nm was read on a Synergy HT microplate reader (BioTek) and measurement was converted to neutrophil number using a standard curve.

### Generation of HXA_3_-containing supernatants to assess the effect of AJC disruption (“infection priming”) of ALI monolayers on PMN migration and bacterial translocation

To assess the role of PLY-induced AJC disruption independent of PLY-induced PMN chemoattractant secretion in barrier function changes upon *Sp* infection, we first generated HXA_3_-containing supernatants by infecting a set of young (2-month-old) or aged (>20-month-old) WT C57BL/6J mouse basal stem cell-derived ALI monolayers with 1×10^7^ WT *Sp* for 1 hour at 37°C with 5% CO_2_, as described (Xu et al., 2024). Infected Transwells were placed into 24-well receiving plates containing HBSS with Ca^2+^ and Mg^2+^ in the apical chamber for an additional 1 hour to allow for polarized chemoattractant secretion. At the end of incubations, apical chamber supernatants, which contain HXA_3_, were collected, centrifuged to remove residual *Sp*, and transferred to new 24 well plates.

To generate ALI monolayers that have been subjected to AJC disruption (“primed”) by infection with *Sp*, we utilized ALI monolayers derived from *Alox15*^*-/-*^ mice, which lack 12-lipoxygenase activity and are incapable of producing HXA_3_. *Sp* grown to log phase were washed and resuspended to 5×10^8^ CFU/ml in HBSS supplemented with Ca^2+^ and Mg^2+^. 25 μl of bacterial suspension was added to the apical surface of the *Alox15*^*-/-*^ ALI monolayers and incubated at 37°C with 5% CO_2_ for 2 hours to allow for priming of the ALI monolayers. These *Sp*-infected *Alox15*^−/−^ ALI monolayers were then placed into plates harboring the HXA_3_-containing supernatants generated above for PMN transmigration and barrier integrity assessment assays.

### Fluorescence microscopy assessment of ALI monolayer integrity

The degree of apical junctional complex integrity and cell confluency of ALI monolayers on Transwell filters was assessed by fluorescence microscopy. Monolayers were fixed in 4% PFA, permeabilized with 0.1% Triton-X 100 in PBS plus 3% BSA, and stained with anti-E-cadherin (24E10, Cell Signaling), followed by Alexa Fluor 488-conjugated anti-rabbit secondary antibody, DAPI, and Alexa Fluor 594-conjugated phalloidin. Transwell filters were then excised and mounted in Vectashield Antifade Mounting Medium (Vector Laboratories) for visualization with a Leica SP8 spectral confocal microscope (Leica). Quantification of E-cadherin junction organization was carried out with the python script IJOQ as previously described (43). Quantitations were normalized to that of untreated controls.

### Neutrophil elastase activity measurements

NE activity in the soluble fraction of BALF from infected mice or PMN supernatants from 1×10^6^ PMNs challenged with 1×10^7^ CFU *Sp* was determined using a PMN Elastase Fluorometric Activity Assay Kit (Abcam), following manufacturer’s instructions. The area under the curve of kinetic substrate conversion curves over 2 hours was measured with a Synergy H1 plate reader (BioTek) and normalized to uninfected controls.

### Presentation of data and statistical analyses

Statistical analysis was carried out using GraphPad Prism (GraphPad Software, San Diego, CA), using ordinary one-way ANOVA followed by Tukey’s *post-hoc* test, or an unpaired t-test in supplemental figures. p values < 0.05 were considered significant in all cases. Tissue and blood bacterial burdens were log-transformed; for all other graphs, the mean values ± SEM are shown. Due to intrinsic donor-to-donor variability of human PMN transmigration efficacy, experiments involving human donors were normalized before pooling individual experiments. The conclusions drawn were those found to be reproducible and statistically significant across independent experiments.

## RESULTS

### Upon pulmonary *Sp* challenge of mice, PLY alters E-cadherin organization prior to promoting PMN influx, epithelial barrier disruption, and bacteremia

Previously, using polarized H292 lung carcinoma cell monolayers, we found that the pore-forming action of PLY triggers a PMN-independent dissolution of E-cadherin (Xu et al., 2023), a junctional protein critical for epithelial barrier function (Bryant & Stow, 2004). To examine the potential role of PLY-mediated *Sp* damage to airway epithelial junctions during pulmonary infection, we followed the kinetics of E-cadherin disorganization, PMN infiltration, barrier disruption, and bacteremia after intratracheal (*i*.*t*.) challenge of 2-month-old BALB/c mice with 1×10^7^ CFU of wild type (WT) or PLY-deficient (Δ*ply*) *Sp*. In accord with our previous results (Xu et al., 2024), the lung burdens of WT- and Δ*ply*-infected mice were indistinguishable during an 18-hour infection, indicating that PLY had no effect on *Sp* fitness in the lungs and that any PLY-associated differences in infection parameters were independent of bacterial load (Figure 1a).

**Figure 1.**
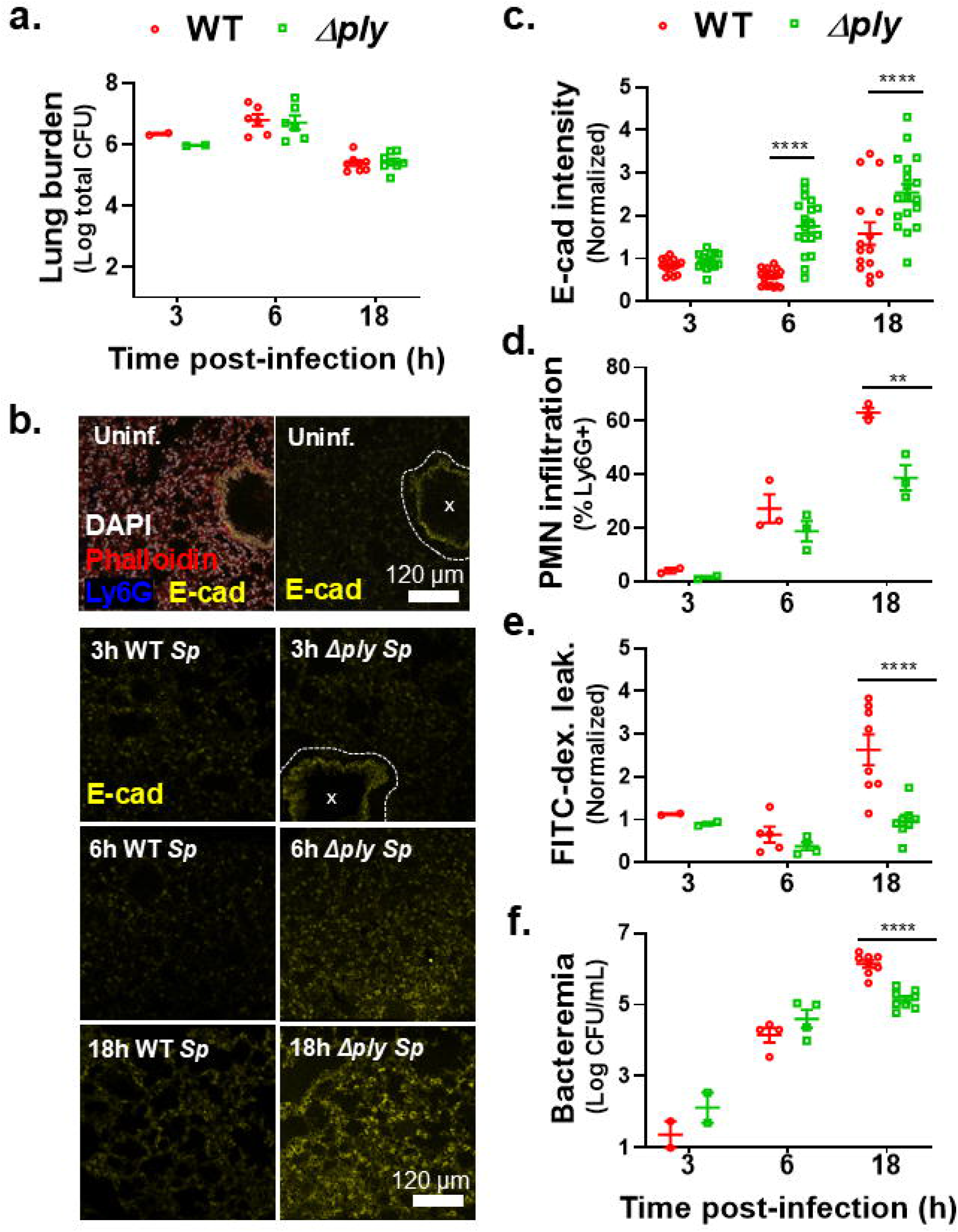
Upon pulmonary *Sp* challenge of mice, PLY alters E-cadherin organization prior to promoting PMN influx, epithelial barrier disruption, and bacteremia. 2-month-old (young) BALB/c mice were infected *i*.*t*. with 1 × 10^7^ wild type (WT) or PLY-deficient mutant (Δ*ply*) *Sp* for 3, 6, or 18 hours. **(a)** Bacterial lung burden determined by measuring CFU in lung homogenates. **(b)** Immunofluorescence (IF) microscopy images of lung sections visualizing E-cadherin (yellow), Ly6G (blue), nuclei (DAPI), and F-actin (phalloidin, red). Bronchial epithelia excluded from analysis are marked by dotted lines. **(c)** Alveolar E-cadherin quantitated by signal intensity analysis in Image J, normalized to uninfected control. **(d)** PMN infiltration determined by flow cytometric enumeration of Ly6G^+^ cells. **(e)** Lung permeability quantitated by measuring the concentration of 70 kDa FITC-dextran in the lung relative to serum after *i*.*v*. administration of FITC-dextran 30 minutes prior to sacrifice, normalized to uninfected control. **(f)** Bacteremia measured by enumerating CFU in whole blood. Each panel is representative of three independent experiments, or pooled data from three independent experiments. Error bars represent mean ± SEM. Statistical analysis between WT and Δ*ply* groups at each time point was performed using ordinary one-way ANOVA with Tukey’s *post-hoc* test: **p-value < 0.01, ****p-value < 0.0001.

To determine a baseline for epithelial junction integrity, we performed immunofluorescence (IF) confocal microscopy of lung sections, visualizing E-cadherin, nuclei, F-actin, and PMNs in uninfected animals. E-cadherin was primarily localized to the bronchial epithelium and a few dispersed and cuboidal (likely alveolar type II) cells along the alveolar epithelium (Figure 1b, “Uninf.”). 3 hours after infection with WT or Δ*ply Sp*, E-cadherin staining patterns (Figure 1b, “3h”) and intensity were unaltered compared to uninfected lung (Figure 1c, “3h”). Flow cytometric quantitation of pulmonary PMNs, identified by the PMN marker Ly6G, revealed that infection by either strain induced no lung inflammation at this time point (Figure 1d). Similarly, epithelial barrier function, assessed by pulmonary leakage of intravenously (*i*.*v*.) administered 70 kDa FITC-dextran, was intact, and few WT or Δ*ply Sp* had yet escaped across the airway barrier into the bloodstream (Figure 1e, f).

Airway epithelial AJCs are upregulated in response to *Sp* invasion, a putative host-protective response (LeMessurier et al., 2013), and at 6 hours post-infection, infection with Δ*ply Sp* resulted in an increase in E-cadherin staining (Figure 1b, c). In contrast, staining after infection with WT *Sp* was 4-fold lower than after infection with Δ*ply Sp* (Figure 1b, c), indicating that PLY disrupted AJC, consistent with our previous observation that PLY promotes E-cadherin dissolution of immortalized cell monolayers (Xu et al., 2023). Regardless of PLY production, PMNs comprised approximately 20% of cells in lung homogenates (Figure 1d) and neither strain caused detectable barrier function compromise at this time point (Figure 1e). Correspondingly, although WT and Δ*ply* were detected in the blood, the level of bacteremia was not influenced by the production of PLY (Figure 1f). Hence, at 6 hours post-infection, PLY production did not alter acute inflammation, barrier function, or bacteremia. Rather, E-cadherin dissolution was the sole PLY-dependent parameter.

Finally, at 18 h post-infection, pulmonary E-cadherin staining of WT *Sp*-infected mice was elevated compared to uninfected mice but remained significantly lower than staining in the lungs of mice infected with Δ*ply Sp*, confirming that PLY promotes E-cadherin disorganization (Figure 1b, c). In agreement with our previous work (Adams et al., 2019; Bhowmick et al., 2013), Δ*ply Sp* triggered less PMN pulmonary infiltration, barrier disruption and bacteremia compared to WT *Sp* 18 h post-infection (Figure 1d-f).

Together, these data reveal that PLY-driven disruption to E-cadherin (AJC) organization precedes PMN influx and epithelial barrier function disturbance.

### PLY-producing *Sp* directly disrupt E-cadherin organization and promote PMN transmigration and barrier disruption independent of concurrent epithelial cell 12-LOX activity

Epithelial monolayers derived from bronchial stem cells and grown with an air-liquid interface (ALI) display many of the architectural and functional attributes of the airway mucosa, including mucus production, cilia, and AJCs that promote a robust junctional barrier (Levardon et al., 2018; Mou et al., 2016; Yonker et al., 2017). We previously utilized apical infection of murine ALI monolayers to recapitulate aspects of *Sp* pathogenesis during mucosal infection (Xu et al., 2024). To investigate PLY-triggered AJC dissolution, we infected human-derived ALI monolayers for 2 hours with 1×10^7^ WT or Δ*ply Sp*, then visualized E-cadherin localization by IF confocal microscopy. In uninfected ALI monolayers, E-cadherin localized to the cell periphery, forming a connected network of circumferential rings (Figure 2a, “Unprimed.”). Upon infection, we observed a loss of this peripheral E-cadherin that was more severe after infection by WT *Sp* than by Δ*ply Sp* (Figure 2a, “WT *Sp*” vs “Δ*ply Sp*”), consistent with our previous work with immortalized cell monolayers (Xu et al., 2023) and the mouse infections described in Figure 1. To quantitate loss of E-cadherin organization, we analyzed images using Intercellular Junction Organization Quotient (IJOQ), which measures the continuity in pericellular junction distribution rather than just signal intensity (Mo et al., 2022). IJOQ quantification indicated that infection by Δ*ply Sp* resulted in a 35% reduction in E-cadherin organization (Figure 2b, “Priming: Δ*ply*”), consistent with the moderate effects of infection of immortalized monolayers by Δ*ply Sp* and indicative of a PLY-independent pathway for AJC disruption (Xu et al., 2023). Infection by PLY-producing (WT) *Sp* resulted in a significantly greater (∼70%) reduction in E-cadherin organization (Figure 2b “Priming: WT”).

**Figure 2.**
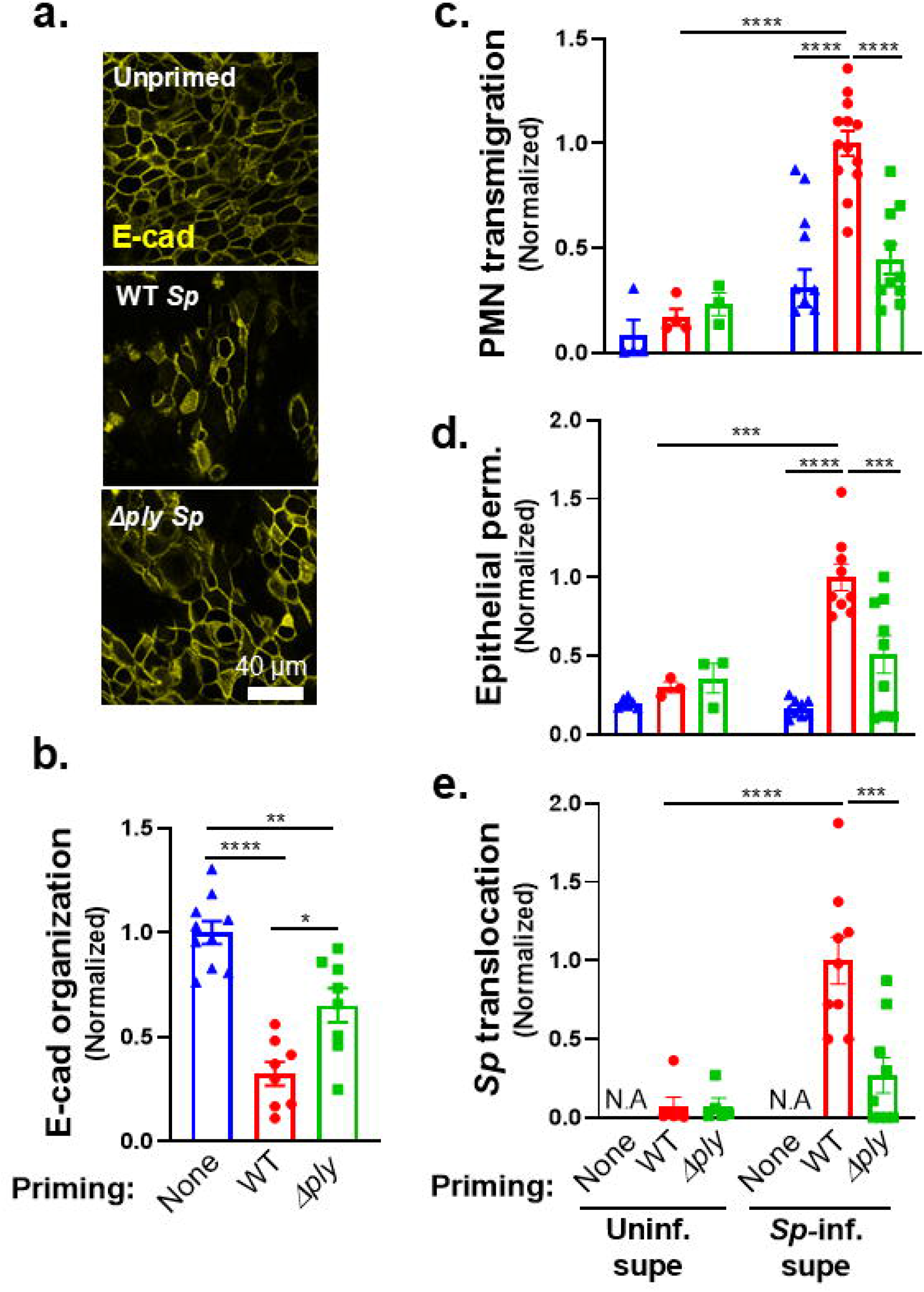
PLY-producing *Sp* directly disrupts E-cadherin organization and promotes PMN transmigration and barrier disruption independent of concurrent epithelial cell 12-lipoxygenase activity. **(a-b)** Healthy adult human BSC-derived ALI epithelial monolayers were infection primed with 1 × 10^7^ WT or Δ*ply Sp*. **(a)** IF microscopy images of fixed and permeabilized monolayers visualizing E-cadherin localization. **(b)** E-cadherin organization quantitated by image analysis via the Intercellular Junction Organization Quotient (IJOQ) script in Python, normalized to no infection priming control. **(c-e)** *Alox15*^*-/-*^ mouse-derived ALI monolayers, which lack 12-lipoxygenase activity and are incapable of generating HXA_3_, were infection primed with 1 × 10^7^ WT or Δ*ply Sp* and transferred into apical chambers containing apical supernatant harvested from uninfected or WT *Sp*-infected WT B6 mouse-derived ALI monolayers. 1 × 10^6^ PMNs were added basally to monolayers and allowed to migrate for 2 hours. Readouts were normalized to WT *Sp*-infection primed ALI monolayers transferred into *Sp*-infection supernatant and include **(c)** the degree of transmigration as determined by MPO activity in the apical chamber, **(d)** epithelial permeability measured by HRP flux, and **(e)** *Sp* translocation quantitated by measuring basolateral CFU. Each panel is representative of three independent experiments, or pooled data from three independent experiments. Error bars represent mean ± SEM. Statistical analyses were performed using ordinary one-way ANOVA with Tukey’s *post-hoc* test: *p-value < 0.05, **p-value < 0.01, ***p-value < 0.001, ****p-value < 0.0001.

The above results indicate that human ALI monolayers, like H292 monolayers (Xu et al., 2023), sustained PMN-independent AJC injury upon infection by PLY-producing *Sp*. This AJC disruption is not sufficient to permit apical to basolateral movement of bacteria in the absence of PMNs (Xu et al., 2024). Nevertheless, sites of AJC disorganization may be exploited as “hotspots” of PMN transmigration, as shown for PMN transendothelial migration (Gronloh, Arts, & van Buul, 2021; Reglero-Real et al., 2021), and we thus investigated whether PLY-triggered AJC dissolution facilitates PMN transmigration and/or compromise of epithelial barrier function. Mouse-derived ALI monolayers, as opposed to those derived from humans, provide an unique, tractable model for examination of this question, as the effects of PLY-induced AJC disruption can be decoupled from PLY-induced PMN chemoattractant secretion by utilizing ALI monolayers derived from *Alox15*^*-/-*^ B6 mice, which are devoid of 12-lipoxygenase activity and incapable of promoting chemotactic PMN movement (Xu et al., 2024). *Alox15*^*-/-*^ ALI monolayers were either infected, i.e., “primed”, with 1×10^7^ WT or Δ*ply Sp* to induce different degrees of AJC disorganization, or as control, left unprimed (i.e., uninfected). To generate HXA_3_-containing medium, we collected supernatant from WT B6 mouse-derived ALI cultures that had been infected with WT *Sp*, then added this (conditioned) “*Sp*-infected” supernatant to the apical chambers of unprimed or primed *Alox15*^*-/-*^ ALI monolayers. Supernatant from uninfected ALI cultures served as a negative control.

1×10^6^ PMNs were added basally and as predicted, control supernatant collected from uninfected ALI monolayers triggered no chemotactic activity, regardless of priming (Figure 2c, “Uninf. supe”). Also, as previously observed (Xu et al., 2024), *Sp*-infected supernatant induced robust PMN transmigration across WT *Sp-*primed *Alox15*^*-/-*^ ALI monolayers (Figure 2c, “WT + *Sp-*inf. supe”). Notably, transmigration across unprimed *Alox15*^*-/-*^ ALI monolayers, which have unperturbed junctions, was minimal and not significantly greater than migration induced by uninfected supernatant (Figure 2c, “None + *Sp*-infected supe”). Similarly, PMN migration across Δ*ply Sp*-primed *Alox15*^*-/-*^ ALI monolayers, which retained a moderate level of E-cadherin organization (Figure 2b, “Priming: Δ*ply*”), was also minimal and 2-fold (and significantly) lower compared to the degree of transmigration across *Alox15*^*-/-*^ ALI monolayers primed with WT *Sp* (Figure 2c, “Δ*ply* + *Sp-*inf. supe”). Given that the above experiments utilized the same *Sp-*infected supernatant to draw PMNs across the monolayers, we conclude that disruption of junctions by PLY-producing *Sp* facilitates subsequent PMN transmigration.

In agreement to the level of PMN transmigration, when we measured barrier disruption by cross-epithelial leakage of the protein marker horseradish peroxidase (HRP) and *Sp* translocation, only WT primed *Alox15*^*-/-*^ ALI monolayers sustained significant HRP flux and *Sp* translocation after migration of PMN to HXA_3_-containing infection supernatant (Figure 2d, e, “WT + *Sp-*inf. supe”). Uninfected *Alox15*^*-/-*^ ALI monolayers remained impermeable to HRP flux after PMN transmigration (Figure 2d, “Uninf. + *Sp-*inf. supe”). Δ*ply-*primed *Alox15*^*-/-*^ ALI monolayers, which sustained less AJC injury and PMN transmigration than WT *Sp* primed ALI monolayers (Figure 2a-c), exhibited lower levels of HRP flux and *Sp* translocation compared to WT (Figure 2d, e, “Δ*ply* + *Sp-*inf. supe”). These data suggest AJCs function as “gate keepers” of PMN influx during *Sp* infection, with PLY-dependent junction disorganization enabling the levels of PMN transmigration that subsequently damage the epithelial barrier and promote *Sp* translocation.

### Aged mice show diminished E-cadherin, increased lung permeability, and increased barrier disruption following pulmonary *Sp* challenge

A decrease in TJ and AJ gene expression occurs in the aging human lung (de Vries et al., 2017), and our findings raise the possibility that age-related changes in airway AJC integrity play a role in promoting susceptibility to *Sp* infection. We performed parallel *i*.*t*. infection of young (2-month-old) or aged (>20-month-old) BALB/c mice with 1×10^7^ CFU of WT or Δ*ply Sp*. Consistent with previous reports (Bhalla et al., 2020), we found that aged mice were highly susceptible to pneumococcal infection. At 18 h.p.i., aged mice infected with WT and Δ*ply Sp* suffered 100-fold and 40-fold higher levels of lung burden than young mice, respectively (Figure 3a). Epithelial AJC integrity assessment by IF at this time point revealed lower levels of alveolar E-cadherin in both WT and Δ*ply* infected aged mice compared to the young (Figure 3b). Image quantification indicated that while young mice exhibited robust E-cadherin upregulation in response to Δ*ply Sp*, this upregulation was diminished more than two-fold by the production of PLY (Figure 3c, “Young” set). In contrast, E-cadherin levels in infected aged mice, regardless of infection by WT or Δ*ply Sp*, were indistinguishable from basal E-cadherin levels of uninfected mice (Figure 3c, “Aged” set).

**Figure 3.**
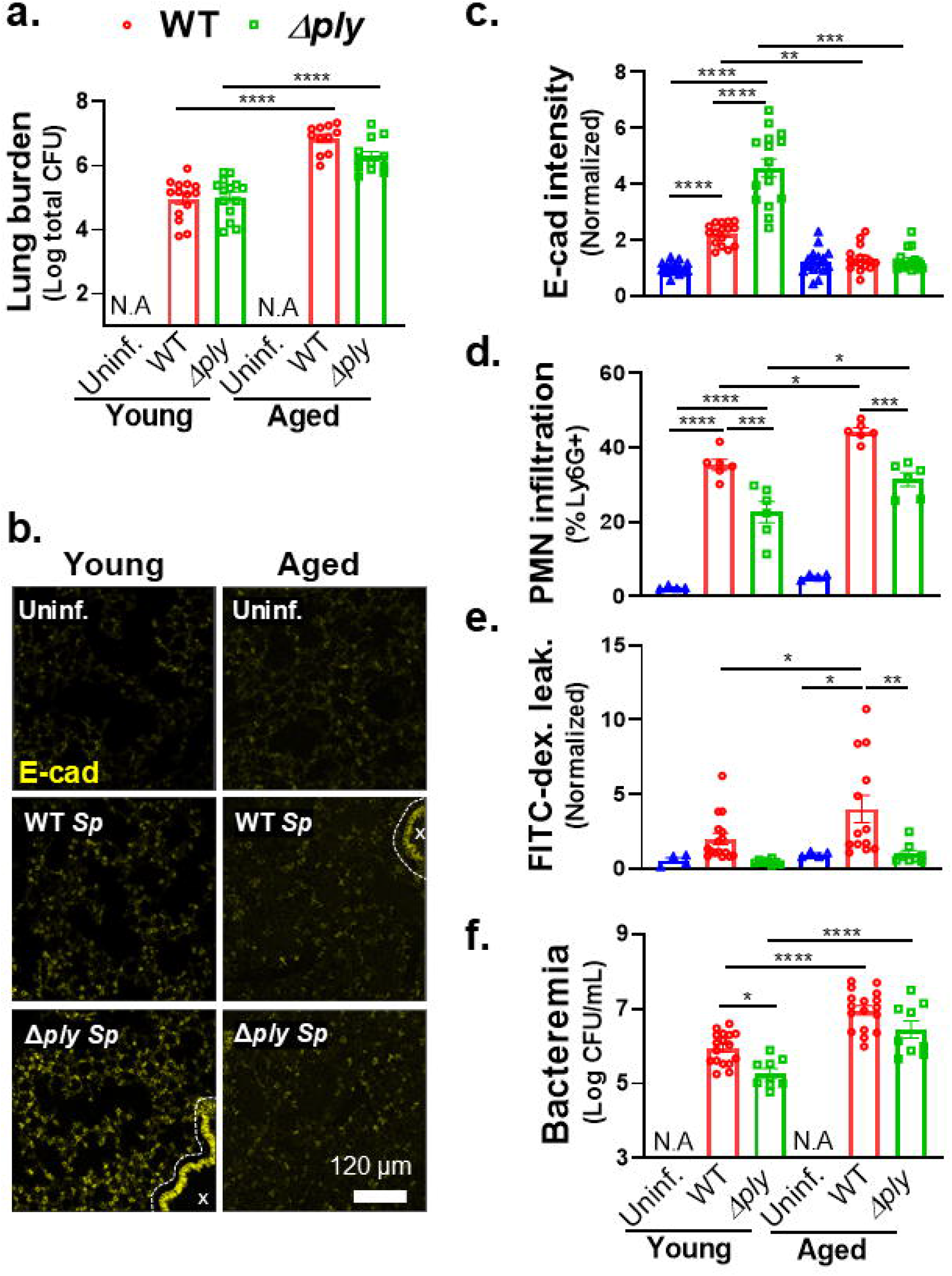
Aged mice show diminished E-cadherin, and increased lung permeability and barrier disruption following pulmonary *Sp* challenge. 2-month-old (young) and 22-month-old (aged) BALB/c mice were infected *i*.*t*. with 1 × 10^7^ WT or Δ*ply Sp* for 18 hours. **(a)** Bacterial lung burden determined by measuring CFU in lung homogenates. **(b)** Lung section IF microscopy images visualizing E-cadherin localization. Bronchial epithelia excluded from analysis are marked by dotted lines. **(c)** Alveolar E-cadherin quantitated by signal intensity analysis in Image J, normalized to uninfected control. **(d)** PMN infiltration determined by flow cytometric enumeration of Ly6G^+^ cells. **(e)** Lung permeability quantitated by measuring the concentration of 70 kDa FITC-dextran in the lung relative to serum after *i*.*v*. administration of FITC-dextran 30 minutes prior to sacrifice, normalized to uninfected control. **(f)** Bacteremia measured by enumerating CFU in whole blood. Each panel is representative of three independent experiments, or pooled data from three independent experiments. Error bars represent mean ± SEM. Statistical analyses were performed using ordinary one-way ANOVA with Tukey’s *post-hoc* test: *p-value < 0.05, **p-value < 0.01, ***p-value < 0.001, ****p-value < 0.0001.

In accord with the lower levels of E-cadherin in aged mice, both WT and Δ*ply Sp-*infected aged mice recruited significantly more PMNs into the airways compared to their respective infections in young mice (Figure 3d). Within each set of (young or aged) infections, WT *Sp* trigger higher PMN infiltration compared to Δ*ply Sp* (Figure 3d), as expected given the previously established role of PLY in PMN recruitment (Adams et al., 2019; Xu et al., 2024). Also, in agreement with the lower levels of E-cadherin and higher levels of PMN infiltration, WT *Sp* infection of aged mice was associated with FITC-dextran leakage that was 5-fold higher than in uninfected aged mice and 2.5-fold higher than in young mice infected with WT *Sp* (Figure 3e, “WT”). Δ*ply Sp* remained unable to inflict substantial damage on the pulmonary barrier despite host age, as no change in FITC-dextran leakage was observed in young or aged mice infected with this strain (Figure 3e, “Δ*ply*”). As expected from the higher lung burden, greater PMN infiltration, and more compromised lung barrier, aged mice suffered higher levels of bacteremia than the young, reflected in 5-fold and 20-fold higher blood burdens upon infection by WT and Δ*ply Sp*, respectively (Figure 3f). Together, these data show that upon *Sp* infection, mice show an age-related decline in AJC integrity and increased PMN infiltration and bacterial spread, with a PLY-dependent increase in pulmonary barrier leakage contributing to the exacerbation of systemic infection.

### Age-related susceptibility to E-cadherin disruption and barrier breach during *Sp* infection is intrinsic to epithelial cells

To investigate whether the age-related AJC defects observed *in vivo* are intrinsic to epithelial cells, we took advantage of the availability of genetically identical ALI monolayers derived from mice of different ages by isolating airway BSCs from young and aged mice to generate corresponding “young” and “aged” ALI epithelial monolayers. All ALI monolayers regardless of age passed the quality control parameter a TEER of at least 1000 Ohms and being impermeable to HRP flux before being used for infection experiments. We found that, as observed above for human ALI monolayers (Figure 2a), E-cadherin was robustly expressed and localized to circumferential rings at cell peripheries in mouse-derived young ALI monolayers (Figure 4a, “Young; Uninf.”). In contrast, E-cadherin staining was weaker and less continuous in aged ALI monolayers, with IJOQ quantification showing a significant 25% reduction in E-cadherin organization compared to young monolayers (Figure 4a, b, “Uninf.”). These data support a cell intrinsic nature of the weakened AJC integrity at baseline that occurs with aging (de Vries et al., 2022).

**Figure 4:**
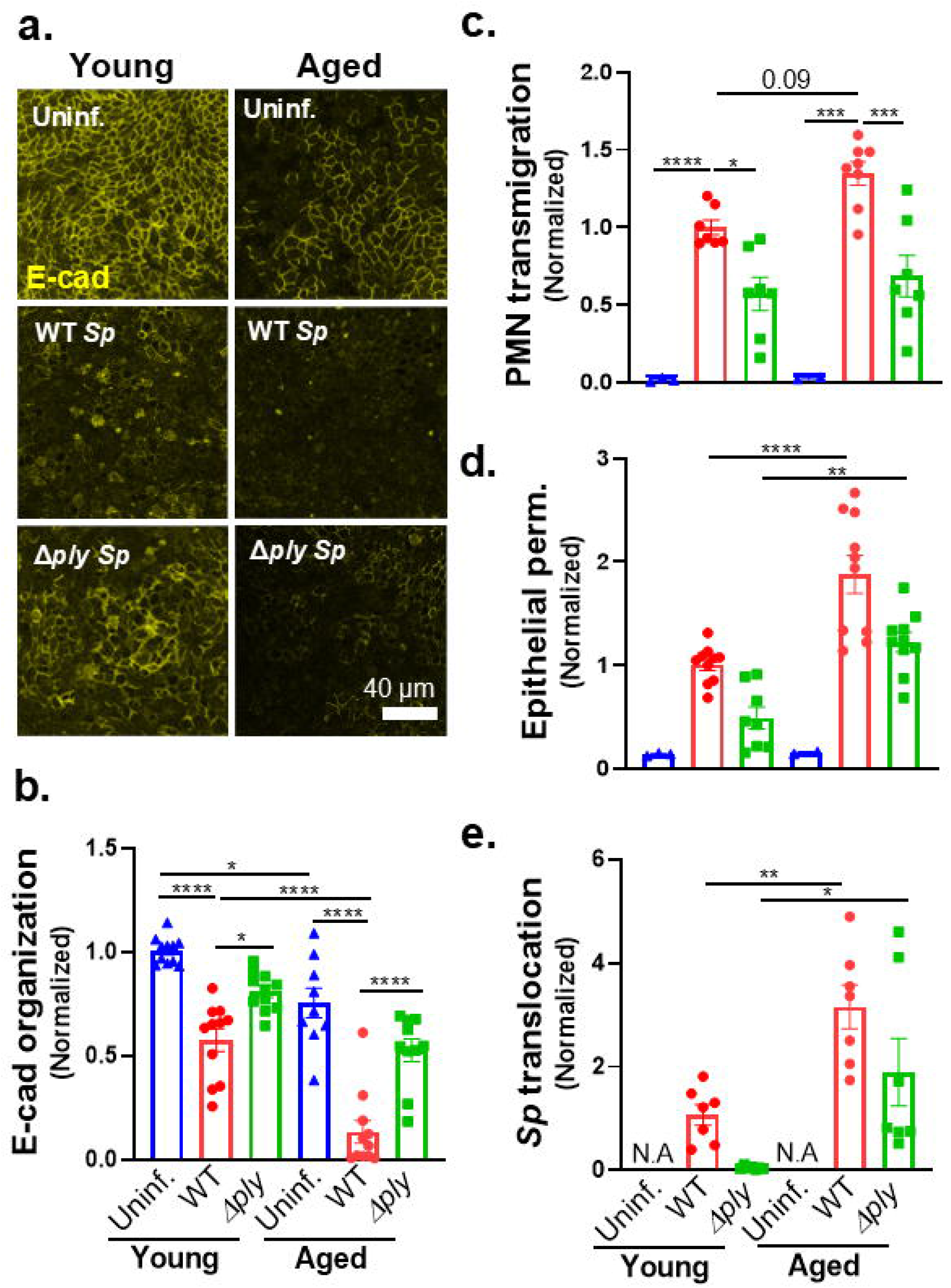
Age-related susceptibility to E-cadherin disruption and barrier breach during *Sp* infection is intrinsic to epithelial cells. 2-month-old (young) and 22-month-old (aged) mouse BSC-derived ALI monolayers were apically infected with 1 × 10^7^ WT or Δ*ply Sp*. **(a)** IF microscopy images of fixed and permeabilized monolayers visualizing E-cadherin localization. **(b)** E-cadherin organization quantitated by image analysis via the IJOQ script in Python, normalized to uninfected young monolayer control. **(c-e)** 1 × 10^6^ PMNs were added basally to monolayers and allowed to migrate for 2 hours. Readouts were normalized to WT *Sp*-infected young monolayer and include **(c)** the degree of transmigration as determined by MPO activity in the apical chamber, **(d)** epithelial permeability measured by HRP flux, and **(e)** *Sp* translocation quantitated by measuring basolateral CFU. Each panel is representative of three independent experiments, or pooled data from three independent experiments. Error bars represent mean ± SEM. Statistical analyses were performed using ordinary one-way ANOVA with Tukey’s *post-hoc* test: *p-value < 0.05, **p-value < 0.01, ***p-value < 0.001, ****p-value < 0.0001.

To assess the effect of aging on the epithelial response to *Sp* infection, we compared the E-cadherin organization of young and aged ALI monolayers after parallel infections with 1×10^7^ WT or Δ*ply Sp*. By imaging and subsequent IJOQ quantitation, infection of young ALI resulted in a loss of E-cadherin organization like that observed above in Figure 2 – infection with WT and Δ*ply Sp* resulted in a 42% and 19% loss of E-cadherin organization, respectively (Figure 4a, b, “Young”). Strikingly, aged ALI monolayers infected with WT *Sp* showed a near complete (87%) dissolution of cell peripheral E-cadherin from an already diminished level of organization (Figure 4a, b, “Aged; WT”); Δ*ply Sp* infection also caused loss of E-cadherin organization but to a lesser extent than WT *Sp* infection (Figure 4a, b, “Aged; Δ*ply*”). These data indicate that AJCs of aged epithelium are both less robust than AJCs of young epithelium and more prone to disruption by *Sp* infection, in accord with the results observed *in vivo* above.

To examine whether the age-associated defect epithelial AJCs enhanced PMN infiltration and/or *Sp* movement across the mucosal barrier, we added young PMNs to the basolateral chamber of *Sp-*infected young or aged ALI monolayers and measured PMN transmigration, HRP flux, and *Sp* translocation. PMNs did not move across uninfected young or aged ALI monolayers (Figure 4c, “Uninf.”). Comparing young and aged ALI monolayers infected with WT *Sp*, aging appeared to result in a slight (1.3-fold; p=0.09) increase in PMN migration in response to WT *Sp* infection and was associated with significant increases in HRP flux (1.8-fold) and *Sp* translocation (3-fold; Figure 4c-e). Although PMN transmigration upon apical infection with Δ*ply Sp* was not significantly different between aged and young ALI (Figure 4c), we found that HRP flux (2.5-fold) and *Sp* translocation (30-fold) were both increased (Figure 4d, e). These age-associated differences were not a result of differential secretion of chemoattactant by *Sp-*infected young and aged ALI monolayers because supernatants collected from the respective monolayers triggered similar levels of PMN transmigration, HRP flux, and *Sp* translocation across *Alox15*^*-/-*^ ALI monolayers (Supplemental Figure 1a-c). Instead, these results suggest that epithelial cell aging, likely the age-associated defect in AJC integrity, contributes to the increased vulnerability of aged compared to young hosts to systemic infection after *Sp* lung inoculation.

### Junction fortification diminishes PMN transmigration, barrier disruption, and bacterial translocation upon *Sp* infection of ALI epithelium derived from young or aged mice

If the age-associated epithelial cell defect in AJC function enhances PMN transmigration, barrier compromise, and bacterial translocation upon apical *Sp* infection, these features should be mitigated by enhancing AJC integrity. Given the importance of E-cadherin for AJC integrity (Nawijn et al., 2011; Yuksel et al., 2021) and its postulated role as a key regulator of age-related barrier defects (de Vries et al., 2022), we treated epithelial monolayers with bardoxolone methyl (CDDO), a Nrf2 agonist that promotes expression of E-cadherin and preserves AJC function in both *in vitro* and in mouse lung injury models (Cheng et al., 2016; Cho & Kleeberger; Ghosh et al., 2022; Guo et al., 2020). Treatment with CDDO indeed enhanced E-cadherin organization in uninfected young mouse ALI monolayers (Figure 5a, b, “Young”: “CDDO” vs “Veh.”) and was associated with resistance to E-cadherin dissolution upon *Sp* infection (Figure 5a, b, “Young”: “*Sp*+CDDO” vs “*Sp*+Veh.”).

**Figure 5:**
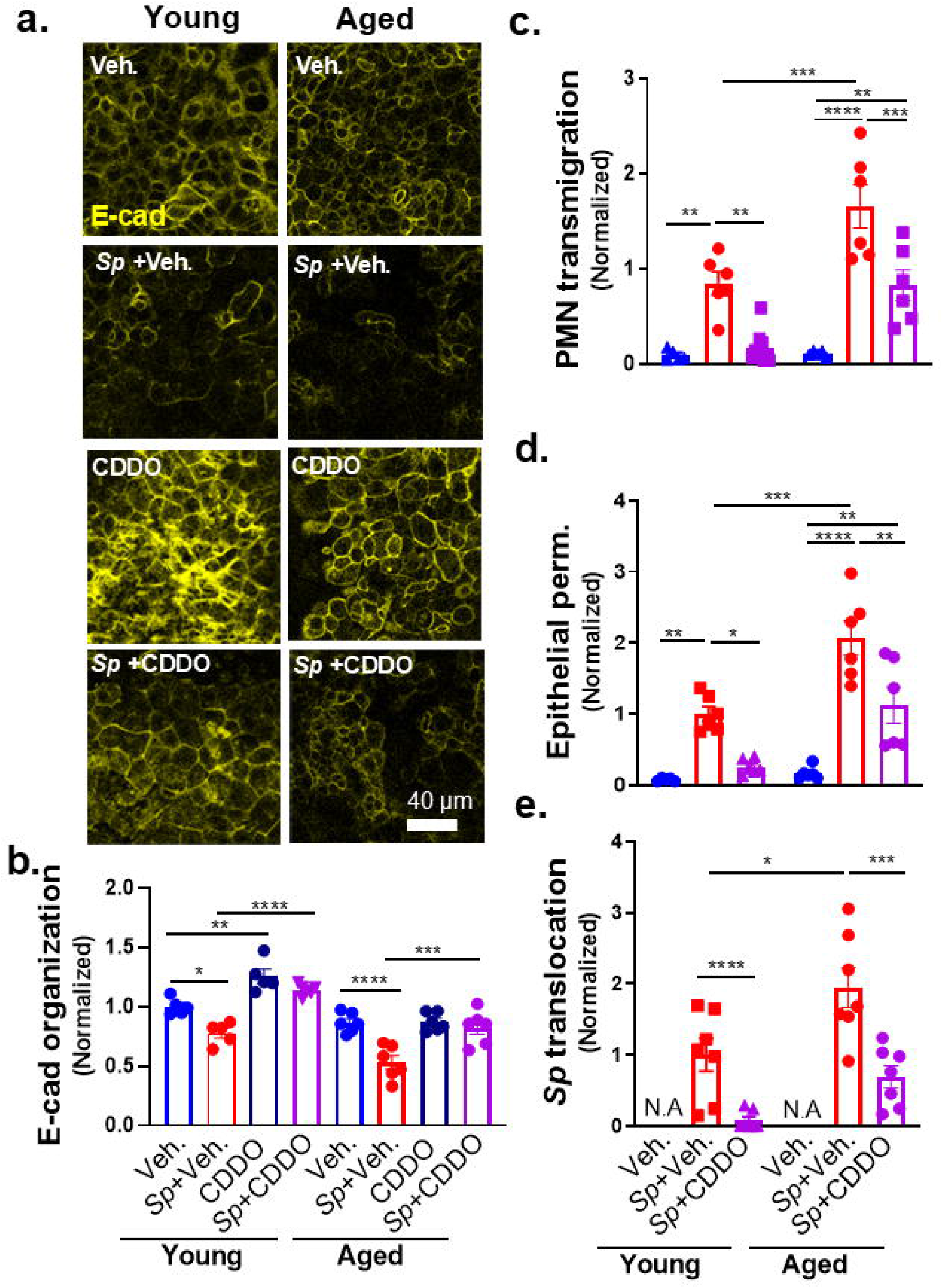
Junction fortification diminishes PMN transmigration, barrier disruption, and bacterial translocation upon *Sp* infection of ALI epithelium derived from young or aged mice. Young and aged mouse BSC-derived ALI monolayers were pretreated with vehicle (DMSO) or 100 nM bardoxolone methyl (CDDO) before apical infection with 1 × 10^7^ WT *Sp*. **(a)** IF microscopy images of fixed and permeabilized monolayers visualizing E-cadherin localization. **(b)** E-cadherin organization quantitated by image analysis via the IJOQ script in Python, normalized to uninfected young monolayer control. **(c-e)** 1 × 10^6^ PMNs were added basally to monolayers and allowed to migrate for 2 hours. Readouts were normalized to WT *Sp*-infected, vehicle-treated young monolayer and include **(c)** the degree of transmigration as determined by MPO activity in the apical chamber, **(d)** epithelial permeability measured by HRP flux, and **(e)** *Sp* translocation quantitated by measuring basolateral CFU. Each panel is representative of three independent experiments, or pooled data from three independent experiments. Error bars represent mean ± SEM. Statistical analyses were performed using ordinary one-way ANOVA with Tukey’s *post-hoc* test: *p-value < 0.05, **p-value < 0.01, ***p-value < 0.001, ****p-value < 0.0001.

Correspondingly, CDDO entirely prevented PMN transmigration, barrier compromise, and bacterial translocation upon *Sp* infection of young ALI monolayers (Figure 5c-e, “Young”: “*Sp*+CDDO” vs “*Sp*+Veh.”).

CDDO pretreatment did not substantially improve E-cadherin organization of uninfected aged ALI monolayers (Figure 5a, b, “Aged”: “CDDO” vs “Veh.”). However, the E-cadherin organization of CDDO-treated aged ALI monolayers, like that of corresponding young ALI monolayers, were protected from *Sp*-mediated disruption (Figure 5a, b, “Aged”: “*Sp*+CDDO” vs “*Sp*+Veh.”). Unlike its effect on young ALI monolayers, CDDO pretreatment of aged ALI monolayers did not entirely abolish PMN transmigration, but mediated a (still significant) 40% reduction (Figure 5c; “Aged”: “*Sp*+CDDO” vs “*Sp*+Veh.”). Correspondingly, CDDO pretreatment of aged ALI monolayers resulted in a 46% and 64% reduction in barrier integrity loss and *Sp* translocation (Figure 5d, e; “Aged”: “*Sp*+CDDO” vs “*Sp*+Veh.”). As predicted, CDDO-mediated AJC fortification of *Alox15*^*-/-*^ ALI monolayers also reduced PMN transmigration, barrier disruption, and *Sp* translocation in response to *Sp*-infected supernatant (Supplemental Figure 2).

### Barrier disruption and *Sp* bacteremia are diminished by junction fortification and NE inhibition in young and aged mice

To test whether the protective effect of CDDO on *Sp*-mediated disruption of AJC in our mouse ALI model corresponded to protection from bacteremia after murine lung infection, we administered CDDO *i*.*p*. to mice one hour before *i*.*t*. challenge. At 18 h.p.i, CDDO pretreatment did not alter bacterial lung burden of either young or aged mice compared to vehicle control (Figure 6a, “*Sp* + CDDO” vs “*Sp* + Veh”), but was assocated with higher E-cadherin levels upon IF microscopy and image quantification of both young and old mice (Figure 6b, c). Notably, CDDO also did not significantly alter PMN infiltration, suggesting that the enhanced E-cadherin did not significantly impede PMN transmigration (Figure 6d, “*Sp* + CDDO” vs “*Sp* + Veh”). Nevertheless, CDDO pretreatment significantly decreased lung permeability to FITC-dextran in young and aged mice by 3-fold and 5-fold, respectively (Figure 6e, “*Sp* + CDDO” vs “*Sp* + Veh”), and, as predicted, diminished bacteremia by 5-fold in the young, and 10-fold in the aged (Figure 6f).

**Figure 6.**
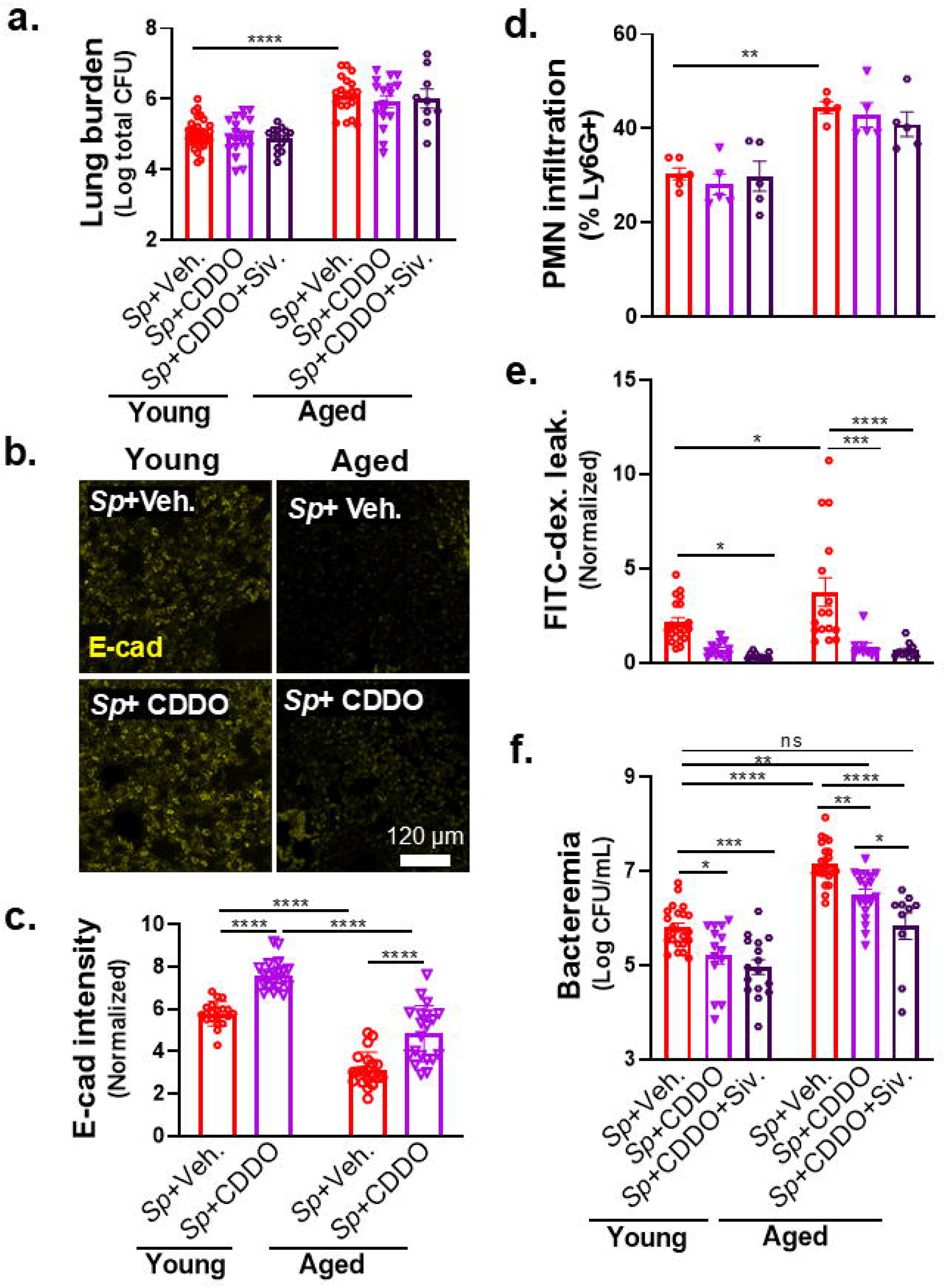
Barrier disruption and *Sp* bacteremia are diminished by junction fortification and NE inhibition in young and aged mice. Young and aged BALB/c mice were treated *i*.*p* with 100 μg/mouse of CDDO or combination of 100 μg/mouse of CDDO and 500 μg/mouse Sivelestat (Siv) 1 hour prior to *i*.*t*. infection with 1 × 10^7^ WT *Sp*. **(a)** Bacterial lung burden determined by measuring CFU in lung homogenates. **(b)** Lung section IF microscopy images visualizing E-cadherin localization. **(c)** Alveolar E-cadherin quantitated by signal intensity analysis in Image J, normalized to uninfected young mouse control. **(d)** PMN infiltration determined by flow cytometric enumeration of Ly6G^+^ cells. **(e)** Lung permeability quantitated by measuring the concentration of 70 kDa FITC-dextran in the lung relative to serum after *i*.*v*. administration of FITC-dextran 30 minutes prior to sacrifice, normalized to uninfected young mouse control. **(f)** Bacteremia measured by enumerating CFU in whole blood. Each panel is representative of three independent experiments, or pooled data from three independent experiments. Error bars represent mean ± SEM. Statistical analyses were performed using ordinary one-way ANOVA with Tukey’s *post-hoc* test: *p-value < 0.05, **p-value < 0.01, ***p-value < 0.001, ****p-value < 0.0001.

Although CDDO pretreatment significantly diminished bacteremia in aged mice, the level of bacteremia, 3.1 × 10^6^ CFU/ml (represented in log_10_ scale in Figure 6f), was still 5-fold higher (p<0.01) than untreated young mice (Figure 6f, “Aged”, “*Sp* + CDDO” vs. “Young”, “*Sp* + veh.”). In our ALI monolayer model, whereas CDDO completely abrogated barrier disruption and bacterial translocation upon *Sp* infection of young ALI monolayers, its effect on these parameters of aged ALI monolayers was only partial (Figure 5d, e, “Aged” vs “Young”; “*Sp* + CDDO” vs “*Sp* + Veh.”), indicating that additional factors may contribute to age-related susceptibility to barrier breach during *Sp* infection. Indeed, E-cadherin is a substrate for NE (Ginzberg et al., 2001; Young, Voisin, Wang, Dangerfield, & Nourshargh, 2007), and we previously showed not only that PMNs from elderly humans exhibit elevated NE activity upon exposure to *Sp* (Bou Ghanem et al., 2017), but also that PMN NE promotes barrier disruption and bacteremia in young mice (Xu et al., 2024). Indeed, here we found that bone marrow-derived PMNs from aged mice, exposed to *Sp*, released higher levels of NE activity compared to bone marrow-derived PMNs from young mice (Supplemental Figure 3a). Correspondingly, upon pulmonary *Sp* infection, higher levels of NE activity were detected in BALF from aged compared to young mice (Supplemental Figure 3b).

The above results indicate that the susceptibility of the aged host to disseminated infection after lung challenge may be due to responses to *Sp* by at least two cell types, i.e., aged PMNs, which secrete higher levels of NE (Bou Ghanem et al., 2017) (Supplemental Figure 3), and aged lung epithelium, which exhibit compromised AJC (Figure 4). We tested whether a therapeutic strategy that addressed both defects would limit bacteremia to levels observed upon infection of untreated young mice. The NE activity inhibitor Sivelestat (Siv) diminishes bacteremia following *Sp* lung challenge in young mice (Xu et al., 2024), so we subjected young and aged mice to dual CDDO and Siv treatment prior to *Sp* pulmonary challenge. In young mice, pulmonary bacterial burden, PMN infiltration, FITC-dextran lung permeability, and bacteremia were unchanged by the addition of Siv to CDDO treatment (Figure 6a, d-f, “Young”: “*Sp* + CDDO + Siv.” vs “*Sp* + CDDO”), suggesting that CDDO and Siv preserve AJCs in a redundant fashion in young hosts. In aged mice, the addition of Siv to CDDO treatment did not alter lung burden or the (already low) FITC-dextran lung permeability associated with CDDO treatment alone (Figure 6a, d-f, “Aged”: “*Sp* + CDDO + Siv.” vs “*Sp* + CDDO”). Dual treatment of aged mice was associated with only a slight (and statistically insignificant) decrease in PMN infiltration (Figure 6d, “Aged”: “*Sp* + CDDO + Siv.” vs “*Sp* + CDDO”). However, the addition of Siv to CDDO treatment of aged mice resulted in an additional 5-fold reduction in bacteremia, which, at 6.7 × 10^5^ CFU/ml, was indistinguishable from the 6.3 × 10^5^ CFU/ml observed for untreated young mice (Figure 6f, “Aged”, “*Sp* + CDDO + Siv.” vs “Young”, “*Sp* + Veh.”). These results indicate that the combination of AJC fortification and NE blockade, by mitigating age-associated, detrimental infection responses of both PMNs and epithelial cells, limits levels of systemic infection to that of a young host.

## DISCUSSION

A vigorous acute inflammatory response is associated with invasive pneumococcal disease (Paterson & Orihuela, 2010; Penaloza et al., 2015; Sundaresh et al., 2021; Yende et al., 2005), so investigation of bacterial and host factors promoting excessive inflammation are likely to provide insight into the root causes of susceptibility to systemic infection by highly vulnerable populations, including the elderly. The major *Sp* virulence factor PLY (Pereira, Xu, Leong, & Sousa, 2022) can enhance pulmonary inflammation by multiple mechanisms (Gonzalez-Juarbe et al., 2015; Parveen et al., 2024; Yoo et al., 2010). We showed that PLY triggers increased epithelial cell production of the lipid chemoattractant HXA_3_, which not only induces PMN influx but also enhances their tissue-destructive capacity, resulting in barrier disruption and bacterial spread (Xu et al., 2023; Xu et al., 2024).

In addition to eliciting damaging inflammatory processes, pathogenic microbes can directly disrupt epithelial barriers to foster systemic spread (Devaux et al., 2019; Gao & Rezaee, 2022; Huber, 2020). Here we showed that during mouse lung infection, PLY promoted AJC disruption independently of its effects on PMN infiltration. Using stem cell-derived ALI epithelial monolayers, which exhibit the cell types and junctional structures of *bona fide* respiratory epithelium, we recapitulated PLY-mediated disruption of E-cadherin organization and barrier function, consistent with our previous finding of PLY-mediated cleavage and mislocalization of E-cadherin in monolayers of immortalized lung epithelial cells (Xu et al., 2023). By employing ALI monolayers deficient in production of HXA_3_ (Xu et al., 2024), we were able to measure the effect of PLY on E-cadherin disorganization independent of its effect on PMN chemoattractant production. Consistent with previous studies of PMN transmigration across epithelial barriers with disrupted AJC (Chun & Prince, 2009; Morton et al., 2016), these experiments revealed that PLY-mediated damage to AJC was required for maximal transmigration of PMNs in response to *Sp*-induced chemoattractant. In fact, PMN transmigration in the absence of PLY-mediated AJC disruption was both minimal and insufficient to significantly diminish barrier function. Finally, fortification of AJC by the Nrf2 agonist CDDO, which increases E-cadherin production, diminished *Sp* translocation in ALI monolayers and bacteremia in lung-inoculated mice.

At baseline and during pneumococcal pneumonia, elderly patients experience elevated PMN pulmonary influx, which correlates with worse clinical outcome (Menter et al., 2014; Pignatti et al., 2011). Aging is accompanied by a heightened state of inflammation, which alters the development of many immune effectors in response to infection (Hinojosa, Boyd, & Orihuela, 2009; Meyer, 2001) and is associated with elevated levels of pro-inflammatory cytokines and chemokines in multiple tissues, including lungs (Bouchlaka et al., 2013; Spencer, Poynter, Im, & Daynes, 1997). In addition, aging is associated with diminished epithelial barrier function (Kling, Lopez-Rodriguez, Pfarrer, Muhlfeld, & Brandenberger, 2017; Thevaranjan et al., 2017). For example, lung barrier function in mice diminishes with age (Kling et al., 2017; Tankersley et al., 2003), and here, comparing the lungs of young or aged mice, we documented an age-associated decline of E-cadherin staining. Furthermore, an increase in E-cadherin production in response to *Sp* lung infection was observed in young but not old mice. Elderly humans exhibit a decrease in expression of AJC genes in the lungs, including that encoding E-cadherin (de Vries et al.; Liu, Zhang, Liang, Noble, & Jiang, 2024), and monolayers derived from bronchial epithelial cells from older individuals exhibit compromised barrier function (de Vries et al.). Utilizing our ALI model of pulmonary epithelium derived from young or aged mice, we found that young and aged epithelium did not differ in their ability to secrete HXA_3_ in response to *Sp* infection; rather, aging was associated with weakened AJCs that were more susceptible to *Sp*-driven AJC damage. Mechano-sensing triggered by PMN migration through endothelial junctions enhances PMN bactericidal activity post-transmigration (Mukhopadhyay et al., 2024) and it is tempting to speculate that the age-associated AJC compromise and the likely concomitant decrease in PMN shear stress may diminish PMN bactericidal capacity. Finally, given that our young and aged ALI monolayers were derived from bronchial stem cells of genetically identical mice, these finding suggest that epigenetic alterations with age, a process implicated in diverse lung diseases (Hagood, 2014), are responsible for the dramatic differences in monolayers derived from these stem cells derived from aged hosts.

The above insights were clearly facilitated by the experimental flexibility provided by mouse ALI monolayers. However, one limitation of the ALI model is that it reflects bronchial, not alveolar epithelium, which comprises most of the pulmonary mucosa and, due to their morphological adaptation to facilitate rapid gas exchange, form AJC comprised of tight junctions alone rather than both tight junctions and adherens junctions as found in bronchial epithelium (Kageyama, Ito, Tanaka, & Nakajima, 2024; Wittekindt, 2017). Unfortunately, *in vitro* studies of *Sp* and PMN interactions with alveolar epithelium awaits the development of a highly physiologic alveolar monolayer model that incorporates PMNs. A second limitation of ALI modeling is that barrier function in the lung involves not just epithelial but also endothelial cells (Kageyama et al., 2024), which are not represented in our ALI monolayers. Nevertheless, PLY disrupts endothelial AJCs (Lucas et al., 2012), suggesting that *Sp* action on endothelium and epithelium may be similar. Furthermore, here we correlated findings derived from our *in vitro* ALI monolayer model with those from our *in vivo* lung infection model. In mice, disruption of the blood-airway barrier, measured by the pulmonary accumulation of intravenously delivered fluorescent dextran or the detection of bacteremia after *i*.*t*. inoculation, reflects translocation across both epithelial and endothelial barriers. Finally, at baseline, the epithelial barrier is approximately ten-fold more stringent than that of endothelium, likely due to the importance of preventing penetration by airway microbes (A. R. Burns, Smith, & Walker, 2003), and the concordance of conclusions from our *in vitro* and *in vivo* models suggests that during *Sp* pneumonia, the epithelium rather than the endothelium may provide the more critical barrier defense.

In addition to age-associated defects intrinsic to epithelium, PMN defects contribute to the risk of systemic *Sp* infections. PMNs from aged mice exhibit a decline in chemotactic accuracy (Sapey et al., 2017) and anti-microbial function (Biasi et al., 1996; Simell et al., 2011), and adoptive transfer of PMNs from young mice into aged mice partially mitigates aged-related susceptibility to *Sp* (Bhalla et al., 2020). In addition to diminished immune defense capabilities, aged PMNs exhibit activities that actively damage tissue (Van Avondt et al., 2023). For example, NE secretion by PMNs plays a key role in barrier disruption and pathogen spread during *Sp* lung infection (Xu et al., 2024), and we found here that not only do PMNs from aged mice secrete increased levels of NE *ex vivo*, but higher levels of this protease were detected in BALF from infected aged mice.

The above findings indicate that both the decline in epithelial AJC integrity and enhanced tissue-destructive capacity of PMNs contribute to the age-associated susceptibility to the disseminated *Sp* infection after *i*.*t*. inoculation. Consistent with this, only by simultaneously targeting both age-related immune defects by administering Sivelestat, which inhibits NE, along with the junction-enhancing agent CDDO, were we able to lower the level of bacteremia in aged mice to that of (untreated) young mice. The requirement for dual treatment to mitigate age-associated risk to disseminated infection underscores the importance of fully characterizing the multifactorial sources of age-associated susceptibility in devising adjunctive therapies to mitigate invasive pneumococcal disease in the elderly.

## ACKNOWLEDGEMENTS

We thank Andrew Camilli for strains and Elsa Bou Ghanem, Jatin M. Vyas, Joan Mecsas, Michael K. Mansour, Byran P. Hurley, and Beth A. McCormick for protocols and helpful discussions. This work was supported by NIH Award AG071268 to JL and JM.

## REFERENCES

Adams, W., Bhowmick, R., Bou Ghanem, E. N., Wade, K., Shchepetov, M., Weiser, J. N., … Leong, J. M. (2019). Pneumolysin Induces 12-Lipoxygenase-Dependent Neutrophil Migration during Streptococcus pneumoniae Infection. J Immunol. doi:10.4049/jimmunol.1800748

Ballinger, M. N., & Standiford, T. J. (2010). Postinfluenza bacterial pneumonia: host defenses gone awry. J Interferon Cytokine Res, 30(9), 643–652. doi:10.1089/jir.2010.0049

Bhalla, M., Simmons, S. R., Abamonte, A., Herring, S. E., Roggensack, S. E., & Bou Ghanem, E. N. (2020). Extracellular adenosine signaling reverses the age-driven decline in the ability of neutrophils to kill Streptococcus pneumoniae. Aging Cell, 19(10), e13218. doi:10.1111/acel.13218

Bhatt, T., Rizvi, A., Batta, S. P., Kataria, S., & Jamora, C. (2013). Signaling and mechanical roles of E-cadherin. Cell Commun Adhes, 20(6), 189–199. doi:10.3109/15419061.2013.854778

Bhowmick, R., Clark, S., Bonventre, J. V., Leong, J. M., & McCormick, B. A. (2017). Cytosolic Phospholipase A2alpha Promotes Pulmonary Inflammation and Systemic Disease during Streptococcus pneumoniae Infection. Infect Immun, 85(11). doi:10.1128/IAI.00280-17

Bhowmick, R., Maung, N., Hurley, B. P., Ghanem, E. B., Gronert, K., McCormick, B. A., & Leong, J. M. (2013). Systemic disease during Streptococcus pneumoniae acute lung infection requires 12-lipoxygenase-dependent inflammation. J Immunol, 191(10), 5115–5123. doi:10.4049/jimmunol.1300522

Biasi, D., Carletto, A., Dell’Agnola, C., Caramaschi, P., Montesanti, F., Zavateri, G., … Bambara, L. M. (1996). Neutrophil migration, oxidative metabolism, and adhesion in elderly and young subjects. Inflammation, 20(6), 673–681. doi:10.1007/BF01488803

Bou Ghanem, E. N., Lee, J. N., Joma, B. H., Meydani, S. N., Leong, J. M., & Panda, A. (2017). The Alpha-Tocopherol Form of Vitamin E Boosts Elastase Activity of Human PMNs and Their Ability to Kill Streptococcus pneumoniae. Front Cell Infect Microbiol, 7, 161. doi:10.3389/fcimb.2017.00161

Bouchlaka, M. N., Sckisel, G. D., Chen, M., Mirsoian, A., Zamora, A. E., Maverakis, E., … Murphy, W. J. (2013). Aging predisposes to acute inflammatory induced pathology after tumor immunotherapy. J Exp Med, 210(11), 2223–2237. doi:10.1084/jem.20131219

Boxio, R., Wartelle, J., Nawrocki-Raby, B., Lagrange, B., Malleret, L., Hirche, T., … Bentaher, A. (2016). Neutrophil elastase cleaves epithelial cadherin in acutely injured lung epithelium. Respir Res, 17(1), 129. doi:10.1186/s12931-016-0449-x

Brown, A. O., Millett, E. R., Quint, J. K., & Orihuela, C. J. (2015). Cardiotoxicity during invasive pneumococcal disease. Am J Respir Crit Care Med, 191(7), 739–745. doi:10.1164/rccm.201411-1951PP

Bryant, D. M., & Stow, J. L. (2004). The ins and outs of E-cadherin trafficking. Trends Cell Biol, 14(8), 427–434. doi:10.1016/j.tcb.2004.07.007

Burns, A. R., Smith, C. W., & Walker, D. C. (2003). Unique structural features that influence neutrophil emigration into the lung. Physiol Rev, 83(2), 309–336. doi:10.1152/physrev.00023.2002

Burns, T., Abadi, M., & Pirofski, L. A. (2005). Modulation of the lung inflammatory response to serotype 8 pneumococcal infection by a human immunoglobulin m monoclonal antibody to serotype 8 capsular polysaccharide. Infect Immun, 73(8), 4530–4538. doi:10.1128/IAI.73.8.4530-4538.2005

Cheng, X., He, S., Yuan, J., Miao, S., Gao, H., Zhang, J., … Wu, P. (2016). Lipoxin A4 attenuates LPS-induced mouse acute lung injury via Nrf2-mediated E-cadherin expression in airway epithelial cells. Free Radic Biol Med, 93, 52–66. doi:10.1016/j.freeradbiomed.2016.01.026

Cho, H. Y., & Kleeberger, S. R. (2015). Association of Nrf2 with airway pathogenesis: lessons learned from genetic mouse models. Arch Toxicol, 89(11), 1931–1957. doi:10.1007/s00204-015-1557-y

Chun, J., & Prince, A. (2009). TLR2-induced calpain cleavage of epithelial junctional proteins facilitates leukocyte transmigration. Cell Host Microbe, 5(1), 47–58. doi:10.1016/j.chom.2008.11.009

Clarke, T. B., Francella, N., Huegel, A., & Weiser, J. N. (2011). Invasive bacterial pathogens exploit TLR-mediated downregulation of tight junction components to facilitate translocation across the epithelium. Cell Host Microbe, 9(5), 404–414. doi:10.1016/j.chom.2011.04.012

GBD LRI Collaborators (2018). Estimates of the global, regional, and national morbidity, mortality, and aetiologies of lower respiratory infections in 195 countries, 1990-2016: a systematic analysis for the Global Burden of Disease Study 2016. Lancet Infect Dis, 18(11), 1191–1210. doi:10.1016/s1473-3099(18)30310-4

Cui, L., Yang, R., Huo, D., Li, L., Qu, X., Wang, J., … Wang, X. (2024). Streptococcus pneumoniae extracellular vesicles aggravate alveolar epithelial barrier disruption via autophagic degradation of OCLN (occludin). Autophagy, 20(7), 1577–1596. doi:10.1080/15548627.2024.2330043

de Vries, M., Faiz, A., Woldhuis, R. R., Postma, D. S., de Jong, T. V., Sin, D. D., … Brandsma, C. A. (2017). Lung tissue gene-expression signature for the ageing lung in COPD. Thorax. doi:10.1136/thoraxjnl-2017-210074

de Vries, M., Nwozor, K. O., Muizer, K., Wisman, M., Timens, W., van den Berge, M., … Brandsma, C. A. (2022). The relation between age and airway epithelial barrier function. Respir Res, 23(1), 43. doi:10.1186/s12931-022-01961-7

Devaux, C. A., Mezouar, S., & Mege, J. L. (2019). The E-Cadherin Cleavage Associated to Pathogenic Bacteria Infections Can Favor Bacterial Invasion and Transmigration, Dysregulation of the Immune Response and Cancer Induction in Humans. Front Microbiol, 10, 2598. doi:10.3389/fmicb.2019.02598

Domon, H., & Terao, Y. (2021). The Role of Neutrophils and Neutrophil Elastase in Pneumococcal Pneumonia. Front Cell Infect Microbiol, 11, 615959. doi:10.3389/fcimb.2021.615959

Frasca, D., & Blomberg, B. B. (2016). Inflammaging decreases adaptive and innate immune responses in mice and humans. Biogerontology, 17(1), 7–19. doi:10.1007/s10522-015-9578-8

Ganesan, S., Comstock, A. T., & Sajjan, U. S. (2013). Barrier function of airway tract epithelium. Tissue Barriers, 1(4), e24997. doi:10.4161/tisb.24997

Gao, N., & Rezaee, F. (2022). Airway Epithelial Cell Junctions as Targets for Pathogens and Antimicrobial Therapy. Pharmaceutics, 14(12). doi:10.3390/pharmaceutics14122619

GBD 2016 Lower Respiratory Infections Collaborators (2018). Estimates of the global, regional, and national morbidity, mortality, and aetiologies of lower respiratory infections in 195 countries, 1990-2016: a systematic analysis for the Global Burden of Disease Study 2016. Lancet Infect Dis, 18(11), 1191–1210. doi:10.1016/s1473-3099(18)30310-4

Ghosh, B., Loube, J., Thapa, S., Ryan, H., Capodanno, E., Chen, D., … Sidhaye, V. K. (2022). Loss of E-cadherin is causal to pathologic changes in chronic lung disease. Commun Biol, 5(1), 1149. doi:10.1038/s42003-022-04150-w

Ginzberg, H. H., Cherapanov, V., Dong, Q., Cantin, A., McCulloch, C. A., Shannon, P. T., & Downey, G. P. (2001). Neutrophil-mediated epithelial injury during transmigration: role of elastase. Am J Physiol Gastrointest Liver Physiol, 281(3), G705–717. doi:10.1152/ajpgi.2001.281.3.G705

Gonzalez-Juarbe, N., Bradley, K. M., Shenoy, A. T., Gilley, R. P., Reyes, L. F., Hinojosa, C. A., … Orihuela, C. J. (2017). Pore-forming toxin-mediated ion dysregulation leads to death receptor-independent necroptosis of lung epithelial cells during bacterial pneumonia. Cell Death Differ, 24(5), 917–928. doi:10.1038/cdd.2017.49

Gonzalez-Juarbe, N., Gilley, R. P., Hinojosa, C. A., Bradley, K. M., Kamei, A., Gao, G., … Orihuela, C. J. (2015). Pore-Forming Toxins Induce Macrophage Necroptosis during Acute Bacterial Pneumonia. PLoS Pathog, 11(12), e1005337. doi:10.1371/journal.ppat.1005337

Gronloh, M. L. B., Arts, J. J. G., & van Buul, J. D. (2021). Neutrophil transendothelial migration hotspots - mechanisms and implications. J Cell Sci, 134(7). doi:10.1242/jcs.255653

Guo, Y., Tu, Y. H., Wu, X., Ji, S., Shen, J. L., Wu, H. M., & Fei, G. H. (2020). ResolvinD1 Protects the Airway Barrier Against Injury Induced by Influenza A Virus Through the Nrf2 Pathway. Front Cell Infect Microbiol, 10, 616475. doi:10.3389/fcimb.2020.616475

Hagood, J. S. (2014). Beyond the genome: epigenetic mechanisms in lung remodeling. Physiology (Bethesda), 29(3), 177–185. doi:10.1152/physiol.00048.2013

Hinojosa, E., Boyd, A. R., & Orihuela, C. J. (2009). Age-associated inflammation and toll-like receptor dysfunction prime the lungs for pneumococcal pneumonia. J Infect Dis, 200(4), 546–554. doi:10.1086/600870

Huber, P. (2020). Targeting of the apical junctional complex by bacterial pathogens. Biochim Biophys Acta Biomembr, 1862(6), 183237. doi:10.1016/j.bbamem.2020.183237

Jacques, L. C., Panagiotou, S., Baltazar, M., Senghore, M., Khandaker, S., Xu, R., … Kadioglu, A. (2020). Increased pathogenicity of pneumococcal serotype 1 is driven by rapid autolysis and release of pneumolysin. Nat Commun, 11(1), 1892. doi:10.1038/s41467-020-15751-6

Kageyama, T., Ito, T., Tanaka, S., & Nakajima, H. (2024). Physiological and immunological barriers in the lung. Semin Immunopathol, 45(4-6), 533-547. doi:10.1007/s00281-024-01003-y

Kim, S., Schein, A. J., & Nadel, J. A. (2005). E-cadherin promotes EGFR-mediated cell differentiation and MUC5AC mucin expression in cultured human airway epithelial cells. Am J Physiol Lung Cell Mol Physiol, 289(6), L1049–1060. doi:10.1152/ajplung.00388.2004

Kling, K. M., Lopez-Rodriguez, E., Pfarrer, C., Muhlfeld, C., & Brandenberger, C. (2017). Aging exacerbates acute lung injury-induced changes of the air-blood barrier, lung function, and inflammation in the mouse. Am J Physiol Lung Cell Mol Physiol, 312(1), L1–L12. doi:10.1152/ajplung.00347.2016

LeMessurier, K. S., Hacker, H., Chi, L., Tuomanen, E., & Redecke, V. (2013). Type I interferon protects against pneumococcal invasive disease by inhibiting bacterial transmigration across the lung. PLoS Pathog, 9(11), e1003727. doi:10.1371/journal.ppat.1003727

Levardon, H., Yonker, L. M., Hurley, B. P., & Mou, H. (2018). Expansion of Airway Basal Cells and Generation of Polarized Epithelium. Bio Protoc, 8(11). doi:10.21769/BioProtoc.2877

Liu, X., Zhang, X., Liang, J., Noble, P. W., & Jiang, D. (2024). Aging-Associated Molecular Changes in Human Alveolar Type I Cells. J Respir Biol Transl Med, 1(3). doi:10.35534/jrbtm.2024.10012

Lucas, R., Yang, G., Gorshkov, B. A., Zemskov, E. A., Sridhar, S., Umapathy, N. S., … Chakraborty, T. (2012). Protein kinase C-alpha and arginase I mediate pneumolysin-induced pulmonary endothelial hyperpermeability. Am J Respir Cell Mol Biol, 47(4), 445–453. doi:10.1165/rcmb.2011-0332OC

Marks, M., Burns, T., Abadi, M., Seyoum, B., Thornton, J., Tuomanen, E., & Pirofski, L. A. (2007). Influence of neutropenia on the course of serotype 8 pneumococcal pneumonia in mice. Infect Immun, 75(4), 1586–1597. doi:10.1128/IAI.01579-06

Menter, T., Giefing-Kroell, C., Grubeck-Loebenstein, B., & Tzankov, A. (2014). Characterization of the inflammatory infiltrate in Streptococcus pneumoniae pneumonia in young and elderly patients. Pathobiology, 81(3), 160–167. doi:10.1159/000360165

Meyer, K. C. (2001). The role of immunity in susceptibility to respiratory infection in the aging lung. Respir Physiol, 128(1), 23–31. doi:10.1016/s0034-5687(01)00261-4

Mo, D., Xu, S., Rosa, J. P., Hasan, S., & Adams, W. (2022). Dynamic Python-Based Method Provides Quantitative Analysis of Intercellular Junction Organization During S. pneumoniae Infection of the Respiratory Epithelium. Front Cell Infect Microbiol, 12, 865528. doi:10.3389/fcimb.2022.865528

Morton, P. E., Hicks, A., Ortiz-Zapater, E., Raghavan, S., Pike, R., Noble, A., … Parsons, M. (2016). TNFalpha promotes CAR-dependent migration of leukocytes across epithelial monolayers. Sci Rep, 6, 26321. doi:10.1038/srep26321

Mou, H., Vinarsky, V., Tata, P. R., Brazauskas, K., Choi, S. H., Crooke, A. K., … Rajagopal, J. (2016). Dual SMAD Signaling Inhibition Enables Long-Term Expansion of Diverse Epithelial Basal Cells. Cell Stem Cell, 19(2), 217–231. doi:10.1016/j.stem.2016.05.012

Mukhopadhyay, A., Tsukasaki, Y., Chan, W. C., Le, J. P., Kwok, M. L., Zhou, J., … Malik, A. B. (2024). trans-Endothelial neutrophil migration activates bactericidal function via Piezo1 mechanosensing. Immunity, 57(1), 52–67 e10. doi:10.1016/j.immuni.2023.11.007

Nawijn, M. C., Hackett, T. L., Postma, D. S., van Oosterhout, A. J., & Heijink, I. H. (2011). E-cadherin: gatekeeper of airway mucosa and allergic sensitization. Trends Immunol, 32(6), 248–255. doi:10.1016/j.it.2011.03.004

Otani, T., & Furuse, M. (2020). Tight Junction Structure and Function Revisited. Trends Cell Biol, 30(10), 805–817. doi:10.1016/j.tcb.2020.08.004

Parveen, S., Bhat, C. V., Sagilkumar, A. C., Aziz, S., Arya, J., Dutta, A., … Subramanian, K. (2024). Bacterial pore-forming toxin pneumolysin drives pathogenicity through host extracellular vesicles released during infection. iScience, 27(8), 110589. doi:10.1016/j.isci.2024.110589

Paterson, G. K., & Orihuela, C. J. (2010). Pneumococci: immunology of the innate host response. Respirology, 15(7), 1057–1063. doi:10.1111/j.1440-1843.2010.01814.x

Penaloza, H. F., Nieto, P. A., Munoz-Durango, N., Salazar-Echegarai, F. J., Torres, J., Parga, M. J., … Bueno, S. M. (2015). Interleukin-10 plays a key role in the modulation of neutrophils recruitment and lung inflammation during infection by Streptococcus pneumoniae. Immunology, 146(1), 100–112. doi:10.1111/imm.12486

Pereira, J. M., Xu, S., Leong, J. M., & Sousa, S. (2022). The Yin and Yang of Pneumolysin During Pneumococcal Infection. Front Immunol, 13, 878244. doi:10.3389/fimmu.2022.878244

Peter, A., Fatykhova, D., Kershaw, O., Gruber, A. D., Rueckert, J., Neudecker, J., … Hippenstiel, S. (2017). Localization and pneumococcal alteration of junction proteins in the human alveolar-capillary compartment. Histochem Cell Biol, 147(6), 707–719. doi:10.1007/s00418-017-1551-y

Pignatti, P., Ragnoli, B., Radaeli, A., Moscato, G., & Malerba, M. (2011). Age-related increase of airway neutrophils in older healthy nonsmoking subjects. Rejuvenation Res, 14(4), 365–370. doi:10.1089/rej.2010.1150

Reglero-Real, N., Perez-Gutierrez, L., Yoshimura, A., Rolas, L., Garrido-Mesa, J., Barkaway, A., … Nourshargh, S. (2021). Autophagy modulates endothelial junctions to restrain neutrophil diapedesis during inflammation. Immunity, 54(9), 1989–2004 e1989. doi:10.1016/j.immuni.2021.07.012

Sapey, E., Patel, J. M., Greenwood, H. L., Walton, G. M., Hazeldine, J., Sadhra, C., … Thickett, D. R. (2017). Pulmonary Infections in the Elderly Lead to Impaired Neutrophil Targeting, Which Is Improved by Simvastatin. Am J Respir Crit Care Med, 196(10), 1325–1336. doi:10.1164/rccm.201704-0814OC

Simell, B., Vuorela, A., Ekstrom, N., Palmu, A., Reunanen, A., Meri, S., … Vakevainen, M. (2011). Aging reduces the functionality of anti-pneumococcal antibodies and the killing of Streptococcus pneumoniae by neutrophil phagocytosis. Vaccine, 29(10), 1929–1934. doi:10.1016/j.vaccine.2010.12.121

Spencer, N. F., Poynter, M. E., Im, S. Y., & Daynes, R. A. (1997). Constitutive activation of NF-kappa B in an animal model of aging. Int Immunol, 9(10), 1581–1588.

Sundaresh, B., Xu, S., Noonan, B., Mansour, M. K., Leong, J. M., & van Opijnen, T. (2021). Host-informed therapies for the treatment of pneumococcal pneumonia. Trends Mol Med, 27(10), 971–989. doi:10.1016/j.molmed.2021.07.008

Tankersley, C. G., Shank, J. A., Flanders, S. E., Soutiere, S. E., Rabold, R., Mitzner, W., & Wagner, E. M. (2003). Changes in lung permeability and lung mechanics accompany homeostatic instability in senescent mice. J Appl Physiol (1985), 95(4), 1681–1687. doi:10.1152/japplphysiol.00190.2003

Thevaranjan, N., Puchta, A., Schulz, C., Naidoo, A., Szamosi, J. C., Verschoor, C. P., … Bowdish, D. M. E. (2017). Age-Associated Microbial Dysbiosis Promotes Intestinal Permeability, Systemic Inflammation, and Macrophage Dysfunction. Cell Host Microbe, 21(4), 455–466 e454. doi:10.1016/j.chom.2017.03.002

Van Avondt, K., Strecker, J. K., Tulotta, C., Minnerup, J., Schulz, C., & Soehnlein, O. (2023). Neutrophils in aging and aging-related pathologies. Immunol Rev, 314(1), 357–375. doi:10.1111/imr.13153

Wenisch, C., Patruta, S., Daxböck, F., Krause, R., & Hörl, W. (2000). Effect of age on human neutrophil function. Journal of Leukocyte Biology. doi:10.1002/jlb.67.1.40

Wittekindt, O. H. (2017). Tight junctions in pulmonary epithelia during lung inflammation. Pflugers Arch, 469(1), 135–147. doi:10.1007/s00424-016-1917-3

Xu, S., Mo, D., Rizvi, F. Z., Rosa, J. P., Ruiz, J., Tan, S., … Adams, W. (2023). Pore-forming activity of S. pneumoniae pneumolysin disrupts the paracellular localization of the epithelial adherens junction protein E-cadherin. Infect Immun, 91(9), e0021323. doi:10.1128/iai.00213-23

Xu, S., Tan, S., Romanos, P., Reedy, J. L., Zhang, Y., Mansour, M. K., … Leong, J. M. (2024). Blocking HXA(3)-mediated neutrophil elastase release during S. pneumoniae lung infection limits pulmonary epithelial barrier disruption and bacteremia. mBio, e0185624. doi:10.1128/mbio.01856-24

Yende, S., Tuomanen, E. I., Wunderink, R., Kanaya, A., Newman, A. B., Harris, T., … Kritchevsky, S. B. (2005). Preinfection systemic inflammatory markers and risk of hospitalization due to pneumonia. Am J Respir Crit Care Med, 172(11), 1440–1446. doi:10.1164/rccm.200506-888OC

Yonker, L. M., Mou, H., Chu, K. K., Pazos, M. A., Leung, H., Cui, D., … Hurley, B. P. (2017). Development of a Primary Human Co-Culture Model of Inflamed Airway Mucosa. Sci Rep, 7(1), 8182. doi:10.1038/s41598-017-08567-w

Yoo, I. H., Shin, H. S., Kim, Y. J., Kim, H. B., Jin, S., & Ha, U. H. (2010). Role of pneumococcal pneumolysin in the induction of an inflammatory response in human epithelial cells. FEMS Immunol Med Microbiol, 60(1), 28–35. doi:10.1111/j.1574-695X.2010.00699.x

Young, R. E., Voisin, M. B., Wang, S., Dangerfield, J., & Nourshargh, S. (2007). Role of neutrophil elastase in LTB4-induced neutrophil transmigration in vivo assessed with a specific inhibitor and neutrophil elastase deficient mice. Br J Pharmacol, 151(5), 628–637. doi:10.1038/sj.bjp.0707267

Yuksel, H., Ocalan, M., & Yilmaz, O. (2021). E-Cadherin: An Important Functional Molecule at Respiratory Barrier Between Defence and Dysfunction. Front Physiol, 12, 720227. doi:10.3389/fphys.2021.720227

