## Supplementary figures and images for "Weakened airway epithelial junctions and enhanced neutrophil elastase release contribute to age-dependent bacteremia risk following pneumococcal pneumonia"

### Supplemental Figure 1

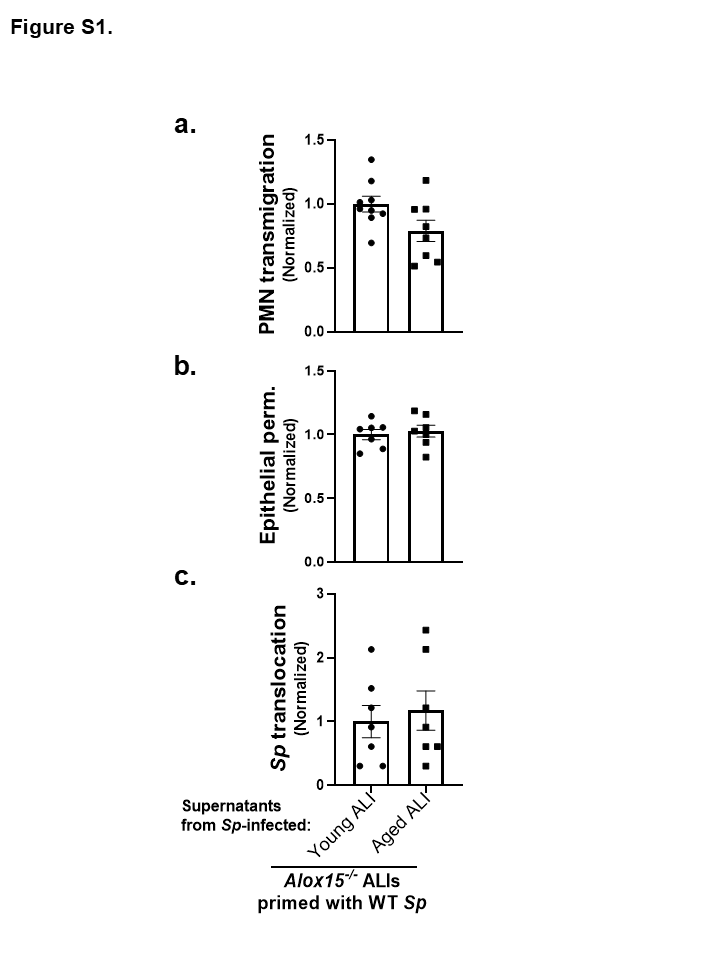

### Supplemental Figure 2

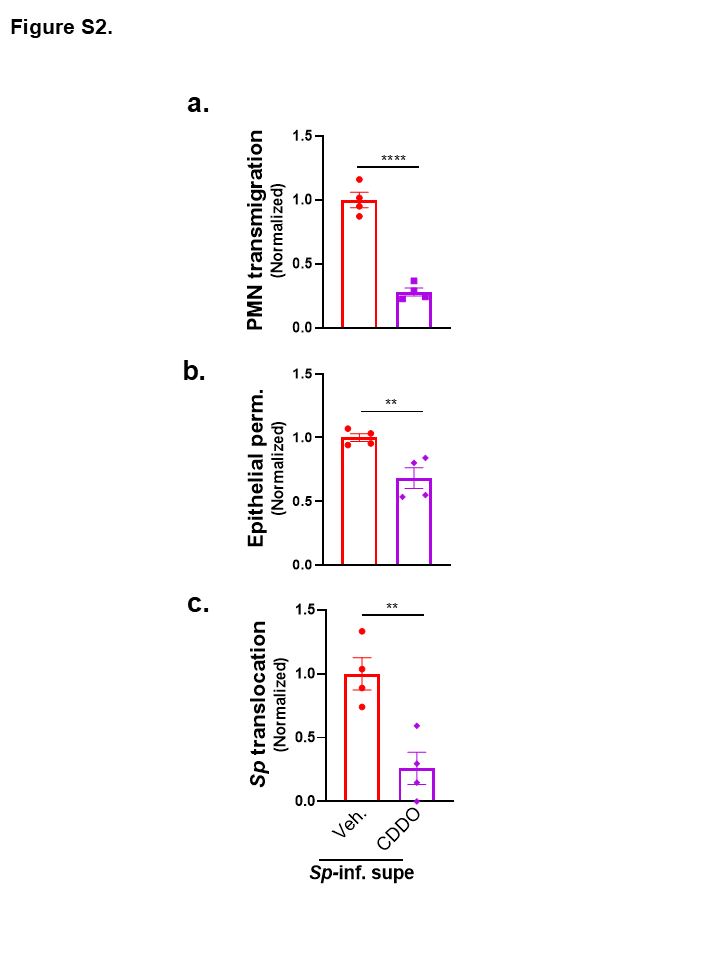

### Supplemental Figure 3

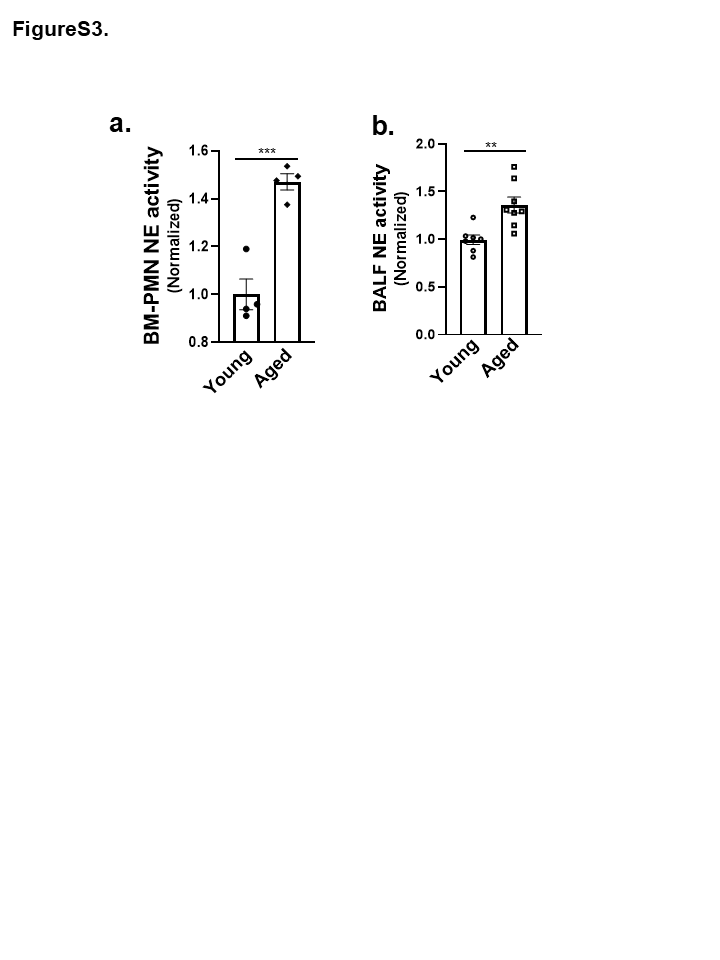
